# Opposing Experiences Shape Parvalbumin Interneuron Plasticity through Distinct Molecular Programs

**DOI:** 10.64898/2026.08.28.747632

**Authors:** Yu Guo, Yu Gao, Sara Knaack, Minjie Shen, Zhiyan Xu, Jiyoun Lee, Yajie Zhang, Paofue Yang, Sabrina X. Huang, Ezra D. Jarzembowski, Drew N. Trygstad, Jensen T. Weik, Jonathan Le, Keegan A. Schoeller, Elena K. Kandror, Abbas H. Rizvi, Xinyu Zhao

## Abstract

Life experiences profoundly influence brain function, yet how opposing experiences are encoded remains unclear. Here, we show that opposing experiences, chronic stress (CS) and voluntary running (VR), induce largely distinct molecular programs rather than a simple bidirectional modulation of a shared molecular program in parvalbumin interneurons (PVIs). PVI-specific translational profiling across frontal cortex and dorsal and ventral hippocampus reveals that experience-dependent gene expression is organized into brain region-specific architectures, which are further structured into distinct co-expression modules. We uncovered CS-induced upregulation of histone acetyltransferase KAT6A and H3K23 acetylation specifically in PVIs of the frontal cortex accompanied by altered genome-wide redistribution of acetylated H3K23, including genes implicated in neuropsychiatric disorders. Targeted elevation of KAT6A in PVIs recapitulates CS-induced morphological and behavioral alterations. Together, our findings reveal that opposing experiences engage distinct and region-specific molecular programs in PVIs and identify a stress-responsive epigenetic pathway underlying inhibitory circuit plasticity.

## Introduction

Psychiatric disorders are complex and heterogeneous conditions affecting a substantial proportion of the global population, with approximately one in seven people worldwide living with a mental disorder^1^. Decades of research have shown that neither genetic background nor environmental exposure alone can fully explain the variability in disease onset, severity, or treatment responses^2^. Gene-environment interactions (G×E) are increasingly recognized as critical determinants of psychiatric vulnerability and are used to explain why individuals with similar genetic backgrounds may exhibit distinct outcomes following negative experiences such as stress and trauma, or positive experiences such as voluntary exercise^3–5^. Early human studies, such as the association between polymorphism of serotonin transporter and severity of stress-induced depression^6^, provided evidence for GxE in mood disorders. More recent human studies and research using animal models have expanded this concept to a wide range of neuropsychiatric disorders including schizophrenia (SCZ), bipolar disorder (BD), and anxiety^7–9^. Importantly, experimental evidence from animal studies has demonstrated that environmental experiences can exert opposing influences on behavioral outcomes: chronic stress (CS) augment anxiety- and depression-like phenotypes, whereas favorable experiences such as exercise buffer stress-related risk, reduce these behaviors, and enhance resilience^10–12^. These contrasting behavioral effects are thought to arise from experience-dependent remodeling of neural circuits^13,14^. However, how the opposing experiences are encoded at the molecular level remains unclear. Specifically, it is unknown whether negative and positive experiences generate opposite molecular states or instead recruit distinct molecular programs within the same neurons.

Parvalbumin-expressing interneurons (PVIs) constitute about 40% of cortical GABAergic interneurons in mice^15^. PVIs are sparsely distributed, fast-spiking cells that provide feedback and feedforward inhibition to excitatory neurons, synchronizing cortical networks and shaping oscillatory activity^15,16^. PVIs have been shown to causally regulate working memory, fear expression, and anxiety-related behaviors, demonstrating their direct influence on cognitive and emotional processing at the circuit level^17–20^. Alterations in PVI number, GABAergic signaling, and gamma oscillations have been reported in psychiatric disorders including SCZ, depression, and anxiety^21–23^. Importantly, PVIs are readily modified by environmental conditions and experience. CS alters glucocorticoid receptor signaling in PVIs and modifies perineuronal nets surrounding PVIs that contributes to excitatory-inhibitory imbalance and behavioral alterations^18,24–27^. In contrast, favorable experiences such as environmental enrichment and voluntary running (VR) remodel PVI structural and network properties in a sustained and reversible manner, reshaping inhibitory circuit states and supporting learning, memory expression, and adaptive emotional regulation^28–30^ and reverses CS-induced reductions in dendritic complexity of PVIs in the medial prefrontal cortex and behavioral impairments^31^. The seemingly opposing effects of positive (e.g., VR) and negative (e.g., CS) experience on PVIs has raised questions whether opposing experience produces simple reciprocal molecular regulation within PVIs or assembles into experience-specific transcriptional architectures shared among brain regions. Only limited work has examined molecular changes within PVIs under stress, typically within a single brain region and without direct comparison to opposing positive experiences^32^. Recent studies have begun to define experience-associated molecular programs in PVIs, including responses to social experience, stress and exercise, using more refined or multimodal approaches^33–36^. However, these efforts remain largely restricted to single experiences or brain regions. Therefore, how opposing environmental experiences organize molecular programs within genetically defined PV interneurons across distinct brain regions remains unresolved.

In this study, we demonstrate that PVIs respond to opposing experiences through largely distinct transcriptional programs rather than simple reciprocal regulation of the same pathways. By profiling PV interneuron translatome across functionally distinct brain regions frontal cortex (FC) and dorsal and ventral hippocampal regions, we reveal that CS and VR engage largely non-overlapping gene networks embedded within regional molecular architectures in a brain-region specific manner. Notably, CS selectively upregulates the histone acetyltransferase KAT6A specifically in FC PVIs, leading to elevated H3K23 acetylation. This is also accompanied by altered genome-wide occupancy by acetylated H3K23, including genes implicated in neuropsychiatric disorders. Remarkably, targeted elevation of KAT6A in PVIs resulted in behavioral deficits similar to those induced by CS. Together, these findings reveal region-embedded molecular programs underlying experience-dependent inhibitory circuit plasticity and identify an epigenetic mechanism linking environmental stress to behavioral changes.

## Results

### Brain region- and experience-specific molecular responses of PVIs to opposing experiences

To investigate how PVIs respond to opposite experiences at gene network levels, we profiled gene expression changes in PVIs in mice that have been subjected to either positive experience (VR) or negative experience (CS). PVIs are sparsely distributed in the brain with a highly branched morphology and extensive connections. To enrich PVI-specific cytosolic mRNAs that are either actively translated or are poised for protein translation, we crossed conditional ribosome tag (Rpl22HA or RiboTag) mice^37^ with PV-Cre driver mice^38^ **(Fig. 1a)**. The resulting PV-HA mice allowed us to identify polyribosome-bound mRNAs specifically in PVIs without cell isolation^35^ ^23,39^. PV-HA mice were subjected to chronic restraint stress (CS), running wheels (VR), or control locked wheels (Ctrl) for 21 days, followed by isolation of the frontal cortex (FC), dorsal hippocampus (dHC) and ventral hippocampus (vHC), the brain regions known to be sensitive to environmental exposure^40,41^, for sequencing analysis **(Fig. 1a)**. These regions represent functionally distinct components of the cognitive-emotion regulation network. The FC integrates sensory and emotional information to support cognitive control and behavioral flexibility^42,43^, the dHC encodes spatial and contextual representations^40^, and the vHC regulates emotional and stress responses through its connectivity with limbic and hypothalamic circuits^40,44^.

**Fig. 1.**
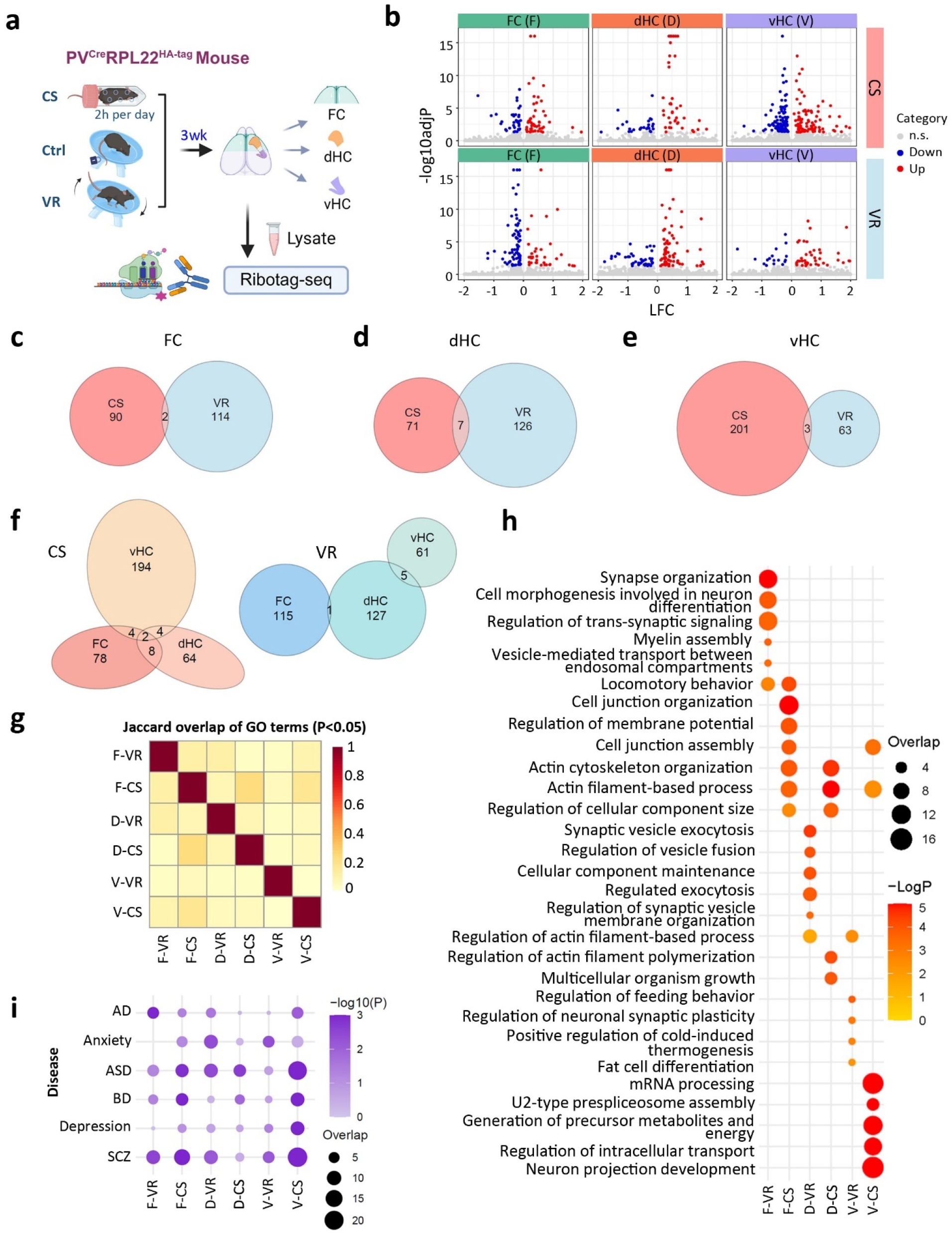
Brain region- and experience-specific transcriptomic responses of parvalbumin interneurons (PVIs) to voluntary running and stress. **a,** Schematic diagram showing experimental design for translational profiling of Parvalbumin interneurons (PVIs) in stress, sedentary and running mice using RiboTag-seq. CS, S: chronic stress; Ctrl, C: control; VR, R: voluntary running. FC, F: Frontal cortex; dHC, D: dorsal hippocampus; vHC, V: ventral hippocampus. **b,** Volcano plots showing differentially translated genes (DTGs) in each region under CS and VR. Genes with adjP < 0.05 were indicated with color (blue, downregulated; red, upregulated). **c-e,** Venn diagram displaying overlap of region-specific DTGs across FC (c), dHC (d), and vHC (e). **f,** Venn diagram displaying overlap of experience-specific DTGs across CS and VR. **g,** Jaccard overlap analysis of all significantly enriched non-redundant GO terms (p < 0.05) across the six conditions (FC/dHC/vHC × CS/VR). The heatmap displays pairwise Jaccard similarity coefficients between DTG-derived GO term sets with hierarchical clustering illustrating region- and experience-dependent functional relationships. **h,** Top five enriched GO Biological Process terms for each DTG set, highlighting distinct functional categories across regions and experiences. The top five most significant terms from each DTG set were selected and the significant enrichments (p<0.05) of all such terms were plotted for across all six DTG sets with the -log10(p) corresponding to color and the number of genes associated with each respective GO term overlap corresponding to size. **i,** Disease enrichment analysis linking region-specific and experience-specific DTGs to neurological and psychiatric disorders. The AD and Anxiety gene set were curated by the DISEASES database. All other gene sets are from DisGeNET. AD: Alzheimer’s disease, ASD: autism spectrum disorder, BD: bipolar disorder, SCZ: schizophrenia. N = 3 mice each condition for Ribotag-seq.

RiboTag immunoprecipitation (IP) of HA-tagged ribosomes together with their associated translating mRNAs, followed by RNA sequencing (RiboTag-seq)^35^ identified differentially translated genes (DTGs) (posterior probability of equivalent expression (PPEE) cutoff < 0.05) (**Fig. 1b; Supplementary Fig. 1a; Supplementary Table 1**). Among the DTGs identified in this analysis, there were minimal overlaps between CS and VR in the same brain region, with only two genes overlapping in FC (**Fig. 1c**), seven in dHC (**Fig. 1d**), and three in vHC (**Fig. 1e**). No DTGs were identified as significantly regulated in opposite directions between CS and VR. Similarly, comparison of DTGs across brain regions under the same condition revealed minimal overlap, with only two genes shared across all three regions in CS, and none in VR **(Fig. 1f).** Together, these findings suggest that PVI responses are both experience-dependent and brain region-specific.

Gene Ontology (GO) analysis of DTGs in each region and condition further revealed distinct pathway signatures **(Fig. 1g, h; Supplementary Fig. 1b, c; Supplementary Table 1)**. Although some GO terms are shared across conditions, Jaccard overlap analysis of all non-redundant GO terms indicated limited functional overlap among the six region-by-experience conditions, reinforcing the conclusion of region- and experience-specific translational remodeling **(Fig. 1g)**. In the FC, DTGs in VR were enriched with synapse-related categories such as synapse organization and trans-synaptic signaling, whereas those in CS were involved cell junction organization, actin cytoskeleton organization, and postsynaptic cytoskeleton structures. In the dHC, VR was enriched for synaptic vesicle exocytosis, regulation of vesicle fusion, and postsynaptic density, while CS showed enrichment in actin cytoskeleton organization, cellular structural remodeling, and chromatin-related binding. In the vHC, VR involved regulation of actin filament-based processes, positive regulation of thermogenesis, and vesicle-mediated transport, whereas CS was enriched for mRNA processing, U2-type prespliceosome assembly, and generation of precursor metabolites and energy, consistent with activation of RNA-processing and metabolic pathways. Overall, the GO analysis highlighted a region- and experience-dependent organization of enriched terms. Across regions, FC and dHC were dominated by cytoskeletal, junctional, and synaptic structural categories, while vHC showed predominant enrichment in RNA-processing and metabolic pathways. Between experiences, VR consistently involved processes related to synaptic organization and vesicle trafficking, whereas CS preferentially enriched categories related to junctional regulation, cytoskeletal organization, transcriptional and RNA-processing pathways, and energy metabolism. These results suggest that experience-dependent transcriptional changes can be grouped into distinct functional programs, which may be further coordinated at the network level.

To further assess the potential disease relevance of these functionally distinct transcriptional programs, we performed disease enrichment analysis using DisGeNET and DISEASE database annotation (**Fig. 1i**). Under CS, the FC showed enrichment in schizophrenia (SCZ), bipolar disorder (BD), and autism spectrum disorder (ASD), while the dHC displayed a more restricted pattern largely centered on ASD-associated gene sets. The vHC exhibited a broader pattern of enrichment, including similar associations as observed in the FC, while additionally extending to Alzheimer’s disease (AD) and other mood-related conditions. In contrast, VR produced a different pattern of disease enrichment, with each region associating with a different subset of disorders. Together, these data highlight the region- and experience-specific disease associations of PVI transcriptional responses.

### Experience-dependent co-expression networks across brain regions

The brain region- and experience-specific transcriptional differences suggest that PVIs adapt to CS and VR through largely distinct but coordinated molecular responses. To investigate how these gene-level changes are organized at the level of co-expression networks, we next applied weighted gene co-expression network analysis (WGCNA) on the combined CS- and VR-DTGs in each region to identify experience-dependent gene modules and their associated functional networks in each brain region **(Fig. 2; Supplementary Fig. 2; Supplementary Table 2)**.

**Fig. 2.**
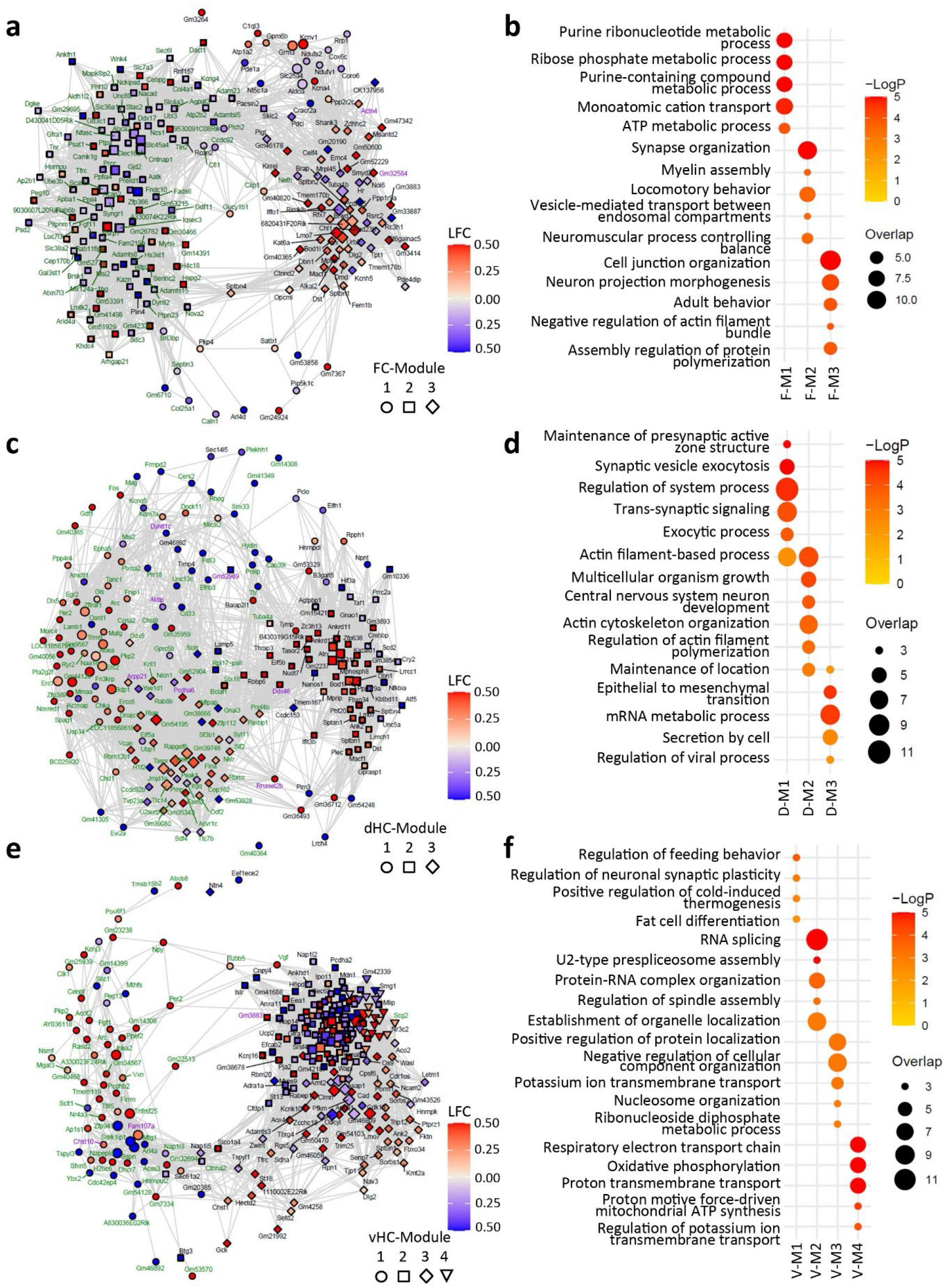
Global co-expression networks across all modules in each region. **a,** Weighted gene co-expression network analysis (WGCNA) of pooled FC DTGs (CS) and VR) identified co-expression modules. **b,** Representative top five GOBP terms are shown for each FC module. **c,** Co-expression modules generated from pooled dHC DTGs. **d,** Representative top five GOBP terms are shown for each dHC module. **e,** Co-expression modules generated from pooled vHC DTGs. **f,** Representative top five GOBP terms are shown for each vHC module. **a**, **c** and **e**: Nodes represent genes, node shapes indicate modules, and label colors represent condition (green: running; black: stress), which naturally segregate in the network. Overlapped genes were shown in purple and showed with LFC under stress condition. Only the top 20% of edges (by weight) are shown to improve visualization clarity, hub genes are highlighted with enlarged shape size. The top five most significant terms for each module DTG set were selected and the significant enrichments (p<0.05) of all such terms were plotted across all modules.

In the FC, three distinct modules represented complementary adaptive processes. Module 1 (circles), composed mainly of CS-associated genes (25 CS, 10 VR), was enriched for pathways related to energy metabolism, transmembrane transport, and synaptic function. Hub genes (large circles) included *Slc25a4* (mitochondrial ADP/ATP carrier) and *Grm3* (metabotropic glutamate receptor 3). Module 2 (squares), dominated by VR genes (103 VR, 5 CS), was associated with synaptic organization and vesicle trafficking-related processes, consistent with enhanced neurotransmission during VR. Its hub gene (large squares) was *Syngr1* (synaptogyrin-1). Module 3 (diamonds), a CS-enriched module (60 CS, 1 VR, 2 shared) was associated with cytoskeletal organization and chromatin-associated processes, with hub genes (large diamonds) including *Tuba1b* (α-tubulin) and *Atrx* (chromatin remodeler) (**Fig. 2a, b; Supplementary Fig. 2a, b**).

In the dHC, three modules reflected experience-dependent regulation of presynaptic and gene regulatory pathways. Module 1 (74 VR, 17 CS, 3 shared) was enriched for presynaptic active-zone assembly and synaptic vesicle exocytosis, consistent with strengthened neurotransmission under VR. Its hub gene *Erc2* (CAST) is a presynaptic active-zone protein. Module 2 (CS-dominant; 53 CS, 4 VR, 2 shared) was associated with cytoskeletal organization and chromatin-associated processes, anchored by *Ppp1r9a* (neurabin-I) and *Dbn1* (drebrin), two actin-associated proteins, while *Atrx* again appeared as a chromatin regulator. Module 3 (VR-associated; 48 VR, 1 CS, 2 shared) was characterized by mRNA processing and secretion-related pathways, with *Srek1*, a splicing regulatory factor, as its hub gene (**Fig. 2c, d; Supplementary Fig. 2c, d**).

In the vHC, four modules illustrated the integration of behavioral, metabolic, and stress-responsive programs. Module 1 (60 VR, 3 CS, 2 shared) was associated with synaptic plasticity and behavioral regulation-related processes, with *Arl4a* and *Fam107a* as hub genes. Module 2 (106 CS, 2 VR, 1 shared) was enriched for RNA splicing-related processes, driven by *Sf3b2* (spliceosome component) and *Numa1* (nuclear mitotic apparatus protein). Module 3 (67 CS) was associated with potassium channel-related activity, protein localization, and synaptic membrane organization, with *Apbb1* (*Fe65*) and *Asap1* as hub genes. Module 4 (triangle) (25 CS, 1 VR) was enriched for mitochondrial oxidative phosphorylation-related pathways, including *Nd1*, *Nd4*, *Cox2*, *Cox3*, and *Atp8* (**Fig. 2e, f; Supplementary Fig. 2e, f**).

Across the three brain regions, VR predominantly engaged synaptic and vesicle trafficking related pathways, whereas CS was associated with postsynaptic, cytoskeletal, and metabolic programs, reflecting structural and epigenetic adaptation. Together, these observations reveal a broad organizational distinction in PVI plasticity, with transmission-related processes preferentially associated with VR and more diverse structural, regulatory, and metabolic programs associated with CS.

### Experience-dependent functional pathway architectures across brain regions

While GO analysis and WGCNA modules revealed region-specific and experience-dependent transcriptional programs and co-expression networks **(Fig. 1 and 2)**, we next asked whether these responses converge into higher-order functional architectures across pathways. To address this, we performed KEGG and Reactome enrichment and organized the results into functional clusters based on shared biological themes.

Under CS, enriched pathways grouped into six major functional clusters **(Fig. 3a, Supplementary Table 3)**: ERK/MAPK signaling cascades, pre-mRNA splicing and RNA processing, mitochondrial oxidative phosphorylation and neurodegeneration-related pathways, secretory pathway and membrane trafficking, neurite outgrowth and cell adhesion-related pathways, and antigen processing and MHC class I presentation. These clusters reflect a combination of signaling, metabolic, structural, and immune-associated processes engaged during CS-associated transcriptional remodeling. VR-associated pathways were organized into five major functional clusters, with limited overlap with the CS-associated pathway network **(Fig. 3b)**: ER-Golgi retrograde trafficking, Rho GTPase signaling pathways, small molecule and ion transport, cell cycle-annotated pathways, and pre-mRNA splicing and RNA processing. Notably, functional clustering revealed largely distinct patterns of organization across brain regions. VR-associated clusters were largely region-specific, whereas CS-associated clusters were more commonly shared across regions, including a mitochondrial oxidative phosphorylation cluster observed across multiple regions.

**Fig. 3.**
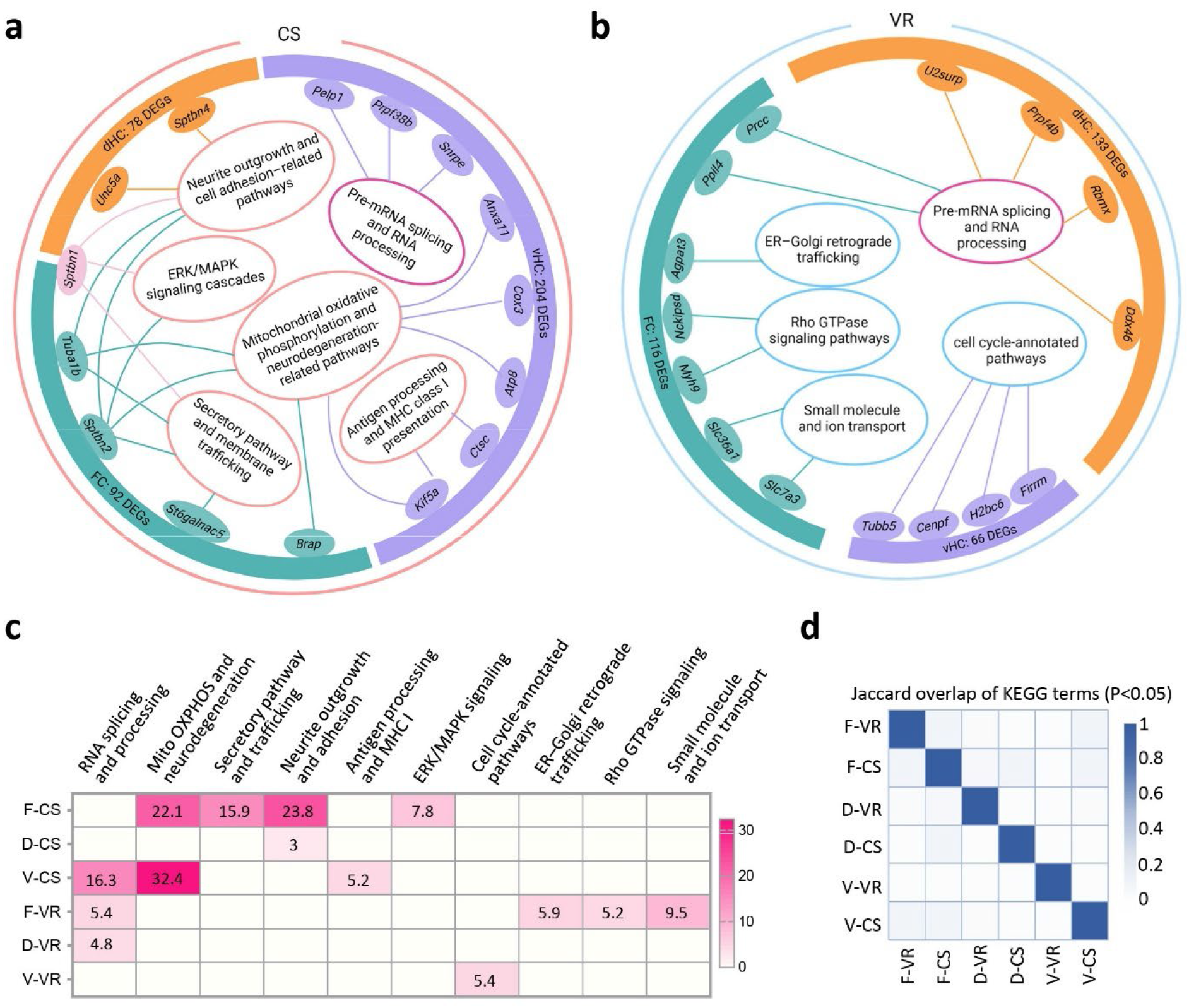
Experience-dependent functional architecture across brain regions. **a,** CS-related architecture. Schematic summary of CS-associated functional changes across FC, dHC, and vHC. Outer layer present brain regions (size indicate the total number of DTGs identified in each region under CS). Inner layer present functional clusters derived from enriched KEGG and Reactome pathways (p < 0.05). Only clusters containing three or more pathway terms were included to construct the functional architecture map. Middle-layer genes represent selected cluster driving genes, defined as DTGs with high fold changes that map to pathways within each functional cluster. **b,** VR-related architecture. Schematic summary for VR-associated functional changes. The network is organized as in (A), enabling direct comparison of functional clusters and their region-specific gene contributors between CS and VR conditions. **c,** Cluster level enrichment summary. Heatmap showing enrichment strength for each functional cluster across region × experience comparisons. For each region and experience, cluster-level enrichment was summarized as the sum total of –log10(p) values across all KEGG and Reactome pathway terms assigned to that functional cluster. **d,** Overlap of enriched KEGG and Reactome pathways. Jaccard overlap analysis comparing enriched pathway terms between conditions and regions.

To quantitatively compare the strength of pathway enrichment across regions and conditions, we summarized enrichment at the cluster level by summing the −log10(P) values of all KEGG and Reactome terms assigned to each functional cluster **(Fig. 3c)**. This analysis revealed distinct regional contributions to each functional program. Jaccard similarity analysis of all enriched KEGG and Reactome pathways revealed minimal overlap across region-condition comparisons **(Fig. 3d)**, indicating that CS and VR largely recruit different sets of molecular pathways even when they converge on similar biological processes. Together, these analyses extend the module-level observations by showing that experience-dependent molecular responses are organized into different higher-order functional architectures across brain regions.

### Cellular context of experience-associated transcriptional programs in the FC

We next performed gene set variation analysis (GSVA) using curated GO terms related to neuronal development, maturation, axonal and dendritic transport, synaptic organization, and behavior (**Supplementary Fig. 3; Supplementary Table 4**). Across all examined pathways, PVIs in the FC exhibited overall higher GSVA enrichment scores compared with those in the dHC and vHC, including axonal transport, synaptic organization, and neuronal morphogenesis, suggesting enhanced engagement of transcriptional programs supporting neuronal structure and connectivity in FC PVIs. Behavior-related terms, including adult behavior, locomotory behavior, and response to stress, also showed stronger positive enrichment in FC PVIs, indicating increased transcriptional engagement in processes linked to behavioral adaptation.

To provide cellular context for the translational programs identified above, we performed multiplexed error-robust fluorescence in situ hybridization (MERFISH) in the FC. UMAP clustering of MERFISH data identified major cell populations, including PVIs, excitatory neurons, astrocytes, oligodendrocytes, and microglia **(Fig. 4a, b)**. Mapping CS- and VR-associated DTGs onto the MERFISH dataset revealed heterogeneous expression patterns of those genes across cell types **(Fig. 4c; Supplementary Table 5)**, with a subset of genes enriched in PVIs and others broadly expressed or preferentially localized to non-PV populations. These findings suggest that many genes identified as CS- or VR-responsive in PVIs are expressed across multiple cortical cell populations rather than being restricted to PVIs.

**Fig. 4.**
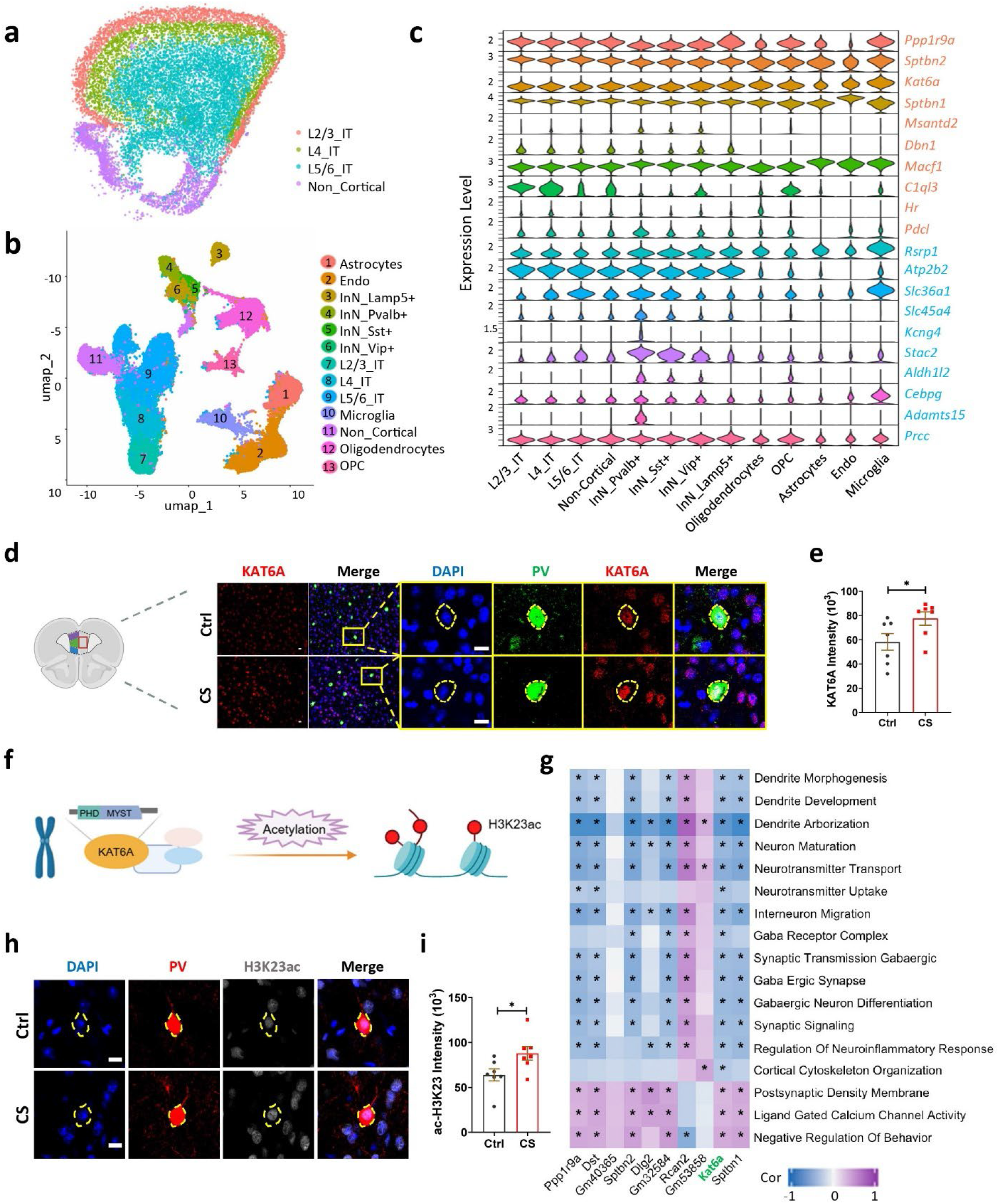
*Kat6a* links neuronal pathways to histone acetylation within FC. **a,** Representative MERFISH field showing spatial localization of cortical layers within FC. **b,** UMAP clustering of MERFISH data from FC showed major cell populations, including PVIs, excitatory neurons, astrocytes, oligodendrocytes, and microglia. **c,** Baseline expression of top FC-CS (orange) and FC-VR (blue) associated DTGs (10 each) across cell types. Genes were selected based on MERFISH probe design constraints and effect size (fold change > 1.2 or < 0.8) and ranked by adjusted p value within each comparison. **d,** Representative confocal images of PVIs in the prefrontal cortex (PFC) of Ctrl or CS mice expressing PV (green) and KAT6A (red). Scale bars: 10 μm. **e,** Immunohistochemistry (IHC) showing increased KAT6A protein expression in the PFC of CS mice compared with controls. **f,** Schematic diagram illustrating KAT6A as a histone acetyltransferase targeting H3K23, leading to the modification of H3K23 acetylation (H3K23ac). **g,** GSVA of FC-CS DTGs across a curated set of phenotypes, including neuronal/synaptic function, cytoskeletal organization, and behavior regulation. Shown are the results for the top 10 most significant DTGs ranked by p value. **h,** Representative confocal images of PVIs in the PFC of Ctrl or CS mice expressing PV (red) and H3K23ac (gray). Scale bars: 10 μm. **i,** IHC of H3K23ac in the PFC, demonstrating elevated acetylation levels consistent with KAT6A upregulation. Data are presented as mean ± s.e.m from N = 7 mice. Two-tailed Student’s test in (**e**) and (**h**). *p < 0.05.

### KAT6A is a key regulator mediating PVI response to CS in FC

To identify potential molecular regulators of PVI translational programs, we focused on mRNAs identified by PVI-RiboTag-seq (**Fig. 1**) across the six region-by-experience conditions. Transcripts with a PPEE < 0.05 and at least a 20% increase or decrease in expression (greater than 1.2 or less than 0.8) were considered candidate regulators. We further narrowed down the candidate by removing transcripts at low levels and considering the availability of antibodies, yielding fifteen candidates for experimental validation using quantitative immunohistology. Among these, protein levels of five candidates (*Chgb, Kat6a, Myh9, Sptbn1, Srek1ip1*) exhibited expression level changes, consistent with the RiboTag-seq results **(Supplementary Fig. 4a)**.

From this validated subset, *Kat6a* (also known as *Moz* or *Myst3*), encoding KAT6A, was selected for further analysis based on its biological relevance and consistency in validation (**Fig. 4d, e**). KAT6A is a histone acetyltransferase that catalyzes histone acetylation, particularly at H3K23 (H3K23ac), modulating chromatin accessibility and transcriptional activation **(Fig. 4f)**^45–48^. *Kat6a* mRNA was widely expressed across FC and in multiple cell types **(Fig. 4c; Supplementary Fig. 4b, c)**. However, in CS mice, KAT6A protein level was significantly increased in PVIs, but not in surrounding non-PV cells in FC **(Supplementary Fig. 4d)**, suggesting that the CS-induced upregulation of KAT6A occurred preferentially within PVIs.

Given that *Kat6a* ranked among the top DTGs in the FC-CS condition, we next examined its functional associations by GSVA with a curated term panel focused on neurodevelopmental and interneuron-related pathways. In this targeted neurofunctional context, *Kat6a* expression showed the most consistent correlations with synaptic signaling, GABAergic synapse, and neuronal maturation pathways **(Fig. 4g; Supplementary Table 6).** These results suggest that *Kat6a* upregulation in FC PVIs is functionally associated with molecular programs supporting interneuron development and circuit plasticity. We next examined whether *Kat6a* upregulation is accompanied by changes in H3K23ac levels in PVIs under FC-CS. IHC analysis revealed that H3K23ac signals were significantly increased **(Fig. 4h, i)**, consistent with the observed upregulation of KAT6A and suggesting enhanced histone acetylation and potential chromatin remodeling in PVIs in response to CS.

### CS-induced chromatin remodeling in FC PVIs engages transcriptional, cellular, and disease-relevant programs

To determine the impact of elevated KAT6A and H3K23ac in PVIs in response to CS, we performed Cleavage Under Targets and Tagmentation (CUT&Tag) sequencing to profile genome wide distribution of H3K23ac in FC PVIs in both CS and control mice. For this experiment, we exposed mice to a modified (6 h/day, 7-day) restraint stress paradigm **(Supplementary Fig. 5a)**, yielding the same total stress exposure (42 h) as the 21-day design (**Fig. 1**) but reduced long-term adaptation and batch variability. KAT6A level was confirmed to be upregulated in FC PVIs in this experimental paradigm **(Supplementary Fig. 5b)**, validating the consistency of our CS models. To enrich nuclei of PVIs, we generated Sun1::PV-Cre mice that had Cre-dependent expression of nuclear-envelope protein Sun1-GFP fusion protein enabling selective labeling of nuclear envelope of PVIs^49^. This system allows for fluorescence-activated sorting of GFP⁺ PVI nuclei from the FC of Ctrl and CS mice for CUT&Tag-seq (**Fig. 5a**). CUT&Tag libraries showed high quality and reproducibility across biological replicates (**Supplementary Fig. 6a, b**). Genome-wide profiling revealed widespread redistribution of H3K23 acetylation under CS, with 2385 hyper-acetylated and 3315 hypo-acetylated peaks identified in FC PVIs (adjP < 0.1; **Fig. 5b-d**; **Supplementary Table 7**). Peak annotation indicated that most signals of H3K23ac localized to intronic and distal intergenic regions (**Fig. 5e**), consistent with previous studies^50^. Motif enrichment analysis of all DA peaks using non-DA H3K23ac peaks as background revealed modest enrichment of several transcription factor motifs, including p53, nuclear receptor, and HMG-box related motifs. However, no dominant motif signature was observed across DA regions (**Supplementary Fig. 6c**), suggesting that stress-associated H3K23ac remodeling may involve multiple regulatory pathways rather than a single predominant transcription factor network. For downstream functional enrichment, only genic peaks (promoter ± 2 kb, UTR, exon, intron, and downstream ≤300 bp) were analyzed to facilitate peak-to-gene assignment and biological interpretation.

**Fig. 5.**
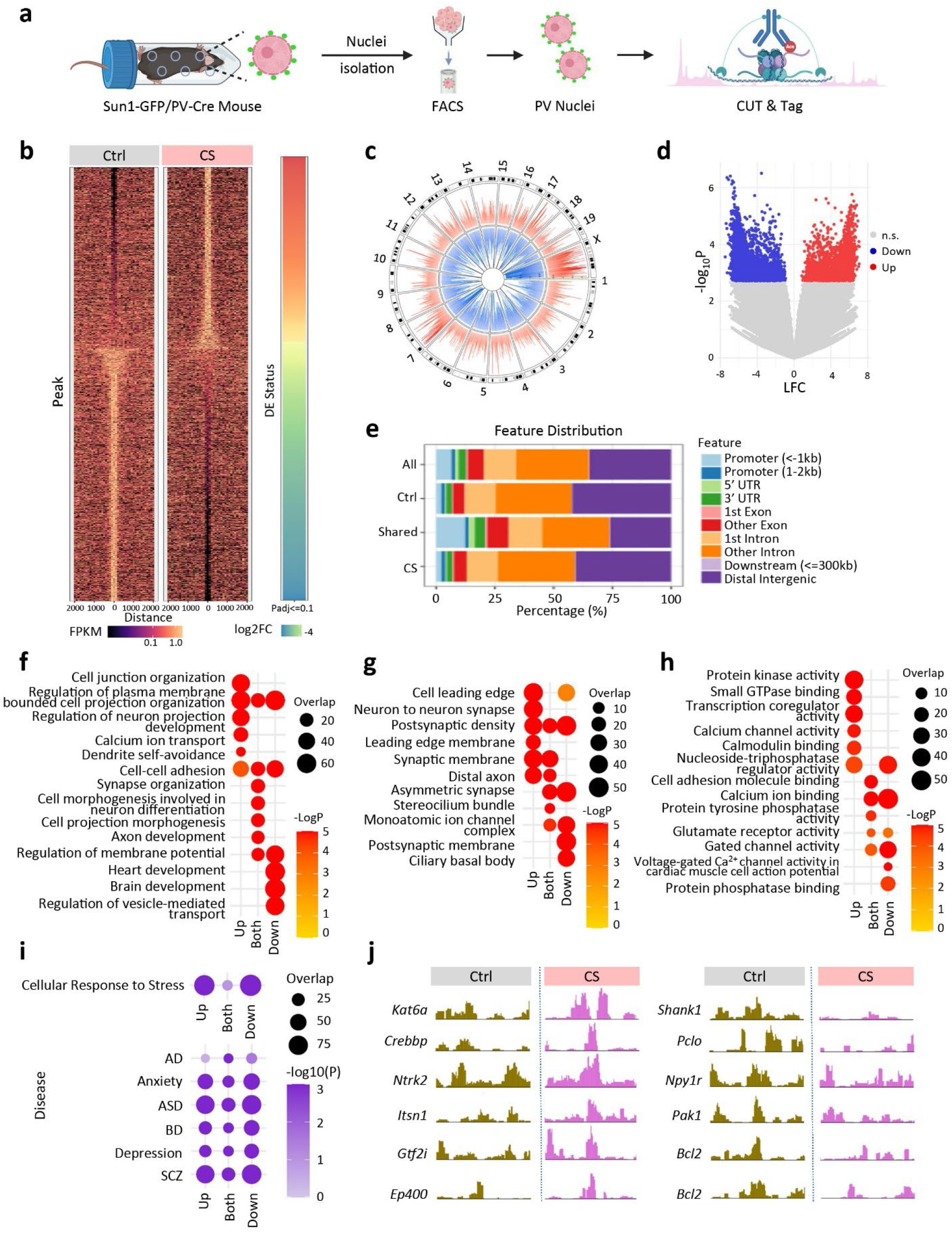
Genome-wide binding profile of H3K23ac in FC of mice subjected to CS. **a,** Schematic diagram illustrating the CUT&Tag workflow. PV interneurons were isolated from Sun1-GFP/PV-Cre mice using flow cytometry, and H3K23ac profiling was performed in CS and Ctrl groups. **b,** Average signal plot displaying genome-wide differences in H3K23ac levels, highlighting regions with increased and decreased acetylation under stress compared with control. **c,** Circos plot showing the chromosomal distribution of H3K23ac signals. The outer ring represents chromosome cytobands. Orange (CS) and blue (Ctrl) tracks indicate H3K23ac signal intensity across the genome, with color gradients reflecting relative signal strength. **d,** Genomic feature distribution across CS-specific, Ctrl-specific, shared, and combined (all) peaks. Most signals localize to intronic and distal intergenic regions. Downstream enrichments used only genic peaks, defined as promoter (±2 kb), Untranslated Regions (UTR), exon, intron, and downstream (≤300 bp), while distal intergenic peaks were excluded. **e,** Volcano plot of differentially acetylated (DA) peaks (with -log10(P) vertical scale), showing significantly enriched and depleted H3K23ac sites. The condition-specificity of DA peaks with Padj value <0.1 are indicated with color (blue, depleted; red, enriched). **f-h,** GO enrichment analysis of genes linked to differentially acetylated (DA) peaks under stress versus control. Only genic peaks (promoter, Untranslated Regions (UTR), exon, intron, and downstream) were considered for gene assignment; distal intergenic peaks were not included. Genes were grouped into Up, Down, and Both based on association with hyper-, hypo-, or mixed acetylation peaks. Top five most significant terms from GOBP (**f**), GOCC (**g**), and GOMF (**h**) for each group were selected and the significant enrichments (p<0.05) of all such terms were plotted for across three groups. **i,** Pathway and disease enrichment analysis of genes linked to CS genic DA peaks. Enriched terms include stress-related signaling pathways and neurological/psychiatric disorders, highlighting the functional relevance of CS-induced H3K23ac changes. **j,** Peak visualization of H3K23ac profiles at representative candidate loci, showing local differences between stress and control.

To determine the cellular processes associated with these acetylation changes, GO enrichment was performed separately for genes associated with H3K23 hyper-acetylated (Up), hypo-acetylated (Down), or both types of peaks (Both) **(Fig. 5f-h; Supplementary Fig. 6d; Supplementary Table 7)**. The analysis revealed both shared and distinct functional signatures across the three groups. Genes associated with hyper-acetylated peaks were enriched for dendritic and synaptic organization, as well as calcium-dependent signaling pathways, suggesting enhanced activity-dependent synaptic remodeling. Genes associated with hypo-acetylated peaks were enriched for synaptic transmission, neuronal signaling, and excitability-related processes, indicating reduced acetylation at genes involved in baseline neuronal function. Genes associated with both hyper- and hypo-acetylated peaks were enriched for processes related to synapse organization, axon development, and regulation of membrane potential, pointing to coordinated regulation of neuronal structure and excitability. Notably, the term related to cellular response to stress was also enriched, although less prominently among genes exhibiting mixed acetylation patterns **(Fig. 5i, top)**, suggesting that these genes may be less involved in canonical stress-response pathways and instead contribute to adaptive modulation of neuronal function. Despite partial overlap in enriched functional categories across the three groups, these gene sets largely converge on shared neuronal processes, such as synaptic organization, neuronal signaling, and excitability. Interestingly, disease enrichment analysis revealed that genes with mixed histone acetylation patterns were more strongly enriched for AD-related gene sets than genes with unidirectional acetylation changes (Up or Down). In contrast, all three groups showed enrichment for gene sets associated with ASD, anxiety, depression, BD, and SCZ, suggesting that associations with these disorders were shared across different patterns of H3K23 acetylation remodeling **(Fig. 5i, bottom)**. Among loci exhibiting CS-associated H3K23ac remodeling, we observed representative genes spanning key functional categories **(Fig. 5j)**, including epigenetic regulators such as *Kat6a*, *Crebbp*, and *Ep400*, activity-dependent signaling molecules including *Ntrk2*, *Pak1*, and *Gtf2i*, genes involved in synaptic organization and vesicle trafficking such as *Shank1*, *Pclo*, and *Itsn1*, and genes involved in stress response and neuronal resilience including *Bcl2* and *Npy1r*. Collectively, these results highlight coordinated epigenetic remodeling at genes linked to neuronal function and CS-related processes.

Together, our findings indicate that CS-associated H3K23ac remodeling in FC preferentially targets genes involved in synaptic organization, neuronal signaling, stress adaptation, and neuropsychiatric disease pathways, consistent with a role for KAT6A-mediated epigenetic remodeling in shaping PVI function under stress.

### Targeted elevation of KAT6A in PVIs impairs neuronal morphology and leads to stress-related behavior deficits

It has been shown that CS leads to reduced dendritic complexity of PVIs^31^ and stress-induced anxiety in mice^51^. We confirmed that PVIs in the medial prefrontal cortex (mPFC) of CS mice indeed have reduced neurite complexities, including fewer intersections decreased total neurite length, fewer branch nodes, and reduced neurite endings compared to control mice **(Supplementary Fig. 7a-e).** Behavioral assessments (**Supplementary Fig. 7f)** showed that CS mice exhibited a significant reduction in time spent in the center zone in the open field test (OF) **(Supplementary Fig. 7g)** and in the open arms in the elevated plus maze test (EPM) **(Supplementary Fig. 7h)**, both indicating increased anxiety-like behavior. In the three-chamber test, CS mice displayed significantly reduced social interest (SI) in another mouse (**Supplementary Fig. 7i**) and reduced preference of a stranger mouse over familiar one (Social Novelty or SN, **Supplementary Fig. 7j**) compared to the control mice.

To determine the significance of elevated KAT6A in PVIs, we induced KAT6A elevation specifically in PVIs, using a CRISPR/Cas9-mediated targeted transcriptional activation (dCas9-Activator or dCas9A) strategy to activate endogenous *Kat6a* gene without excessive overexpression artifacts **(Fig. 6a)**^52^. Five sgRNAs were designed to target the proximal promoter (−234 bp to +165bp relative to transcription start site, TSS) of the mouse *Kat6a* gene. Among them, sgRNA candidates #5 exhibited strong effects in elevating the endogenous *Kat6a* mRNA levels (2-fold) in transfected Neuro2A cells (**Fig. 6b)**. We then cloned *Kat6a* sgRNA #5 into an adeno associate viral (AAV) vector (AAV8) to generate recombinant AAV-*sgKat6a-*mCherry (AAV8-*sgKat6a*-hSyn1-flex-mCherry) virus and stereotaxically injected AAV-*sgKat6a* into the medial PFC of adult male PV-Cre::dCas9-Activator double transgenic (KAT6A-EE) or single transgenic controls (**Fig. 6a**). Immunostaining confirmed approximately 1.5-fold higher KAT6A protein levels in PVIs of KAT6A-EE mice compared with controls **(Fig. 6c, d)**, comparable to the increase observed in PVIs under CS (**Fig. 4e; Supplementary Fig. 5b**). Morphological analysis of mCherry-labeled PVIs showed significant reductions in neurite complexity in KAT6A-EE mice **(Fig. 6e-i)**, recapitulating the changes observed in CS wild-type mice.

**Fig. 6.**
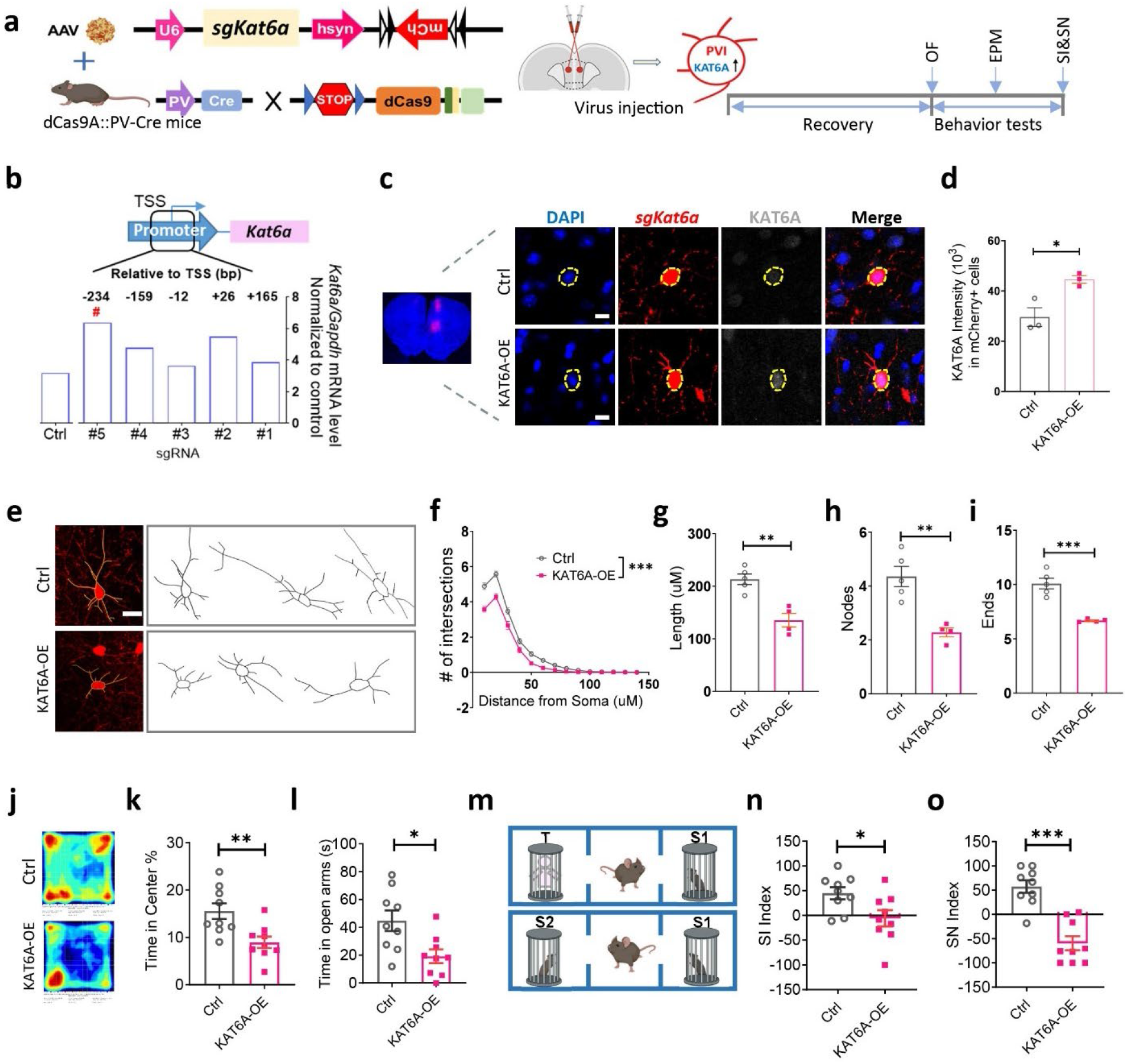
KAT6A upregulation impairs PVI morphology and induces behavioral deficits resembling those in CS. **a,** Experimental strategy for assessing cognitive functions of targeted *Kat6a* gene activation in PVIs of the PFC. Activation of endogenous *Kat6a* gene in the PFC was achieved through stereotaxic injection of AAV expressing *Kat6a*-targeting guide RNA (AAV8-sg*Kat6a*-hSyn1-flex-mCherry) into the medial PFC of PV-Cre/dCas9Activator double transgenic mice or single transgenic mice as control. **b,** Quantification of mRNA levels of *Kat6a* in the Neuro2A cells transfected with *sgKat6a* (or *sgCtrl*) with dCas9-VP64 and SAM. Five sgRNAs were designed to target the proximal promoter region (+165 to -234 bp from the TSS) of *Kat6a*. # indicate the sgRNA used in subsequent experiments. **c,** Representative confocal images of neurons in the prefrontal cortex (PFC), *sgKat6a*-mCherry (red), KAT6A (gray) in AAV-injected PV-Cre/dCas9Activator mice, assessed after behavioral tests. Scale bars: 10 μm. **d,** Quantification of KAT6A intensity in mCherry+ neurons in mPFC. **e,** Representative confocal images and traces of mCherry+ neurons for assessing the effect of KAT6A upregulation on PVI morphology. Scale bar, 20 μm. **f,** Sholl analysis of KAT6A-EE (LV-*sgKat6a, het/het*) and control (LV-*sgKat6a*, wt/het) PVIs. **g-i,** Total neurite length (**g**), total number of nodes (**h**) and total neurite endings (**i**) of KAT6A-EE and control PVIs. **j,** Heatmaps of open field exploration. **k,** Anxiety levels assessed by open field (OF) test. **l,** Anxiety levels assessed by EPM test. **m,** Experimental scheme of testing social interest (SI) and social novelty (SN). **n,o,** KAT6A-EE mice exhibited reduced social interest (SI, **n**), and social recognition (SN, **o**). Data are presented as mean ± s.e.m. Two-tailed Student’s test in (**d**), (**g-i**), (**k-l**) and (**n-o**) from N ≥ 3 mice. MANOVA in (**f**), *F*(1,58) = 35.728, *p* < 0.001, n = 75 cells from 5 mice in wt/het, n = 60 cells from 4 mice in het/het. *p < 0.05, **p < 0.01, ***p < 0.001.

KAT6A-EE mice showed increased anxiety-like phenotypes in both OF **(Fig. 6j, k)** and EPM tests **(Fig. 6l)** and showed reduced social interest and social novelty in the three-chamber tests **(Fig. 6m-o)**, closely resembling the behavioral pattern of CS mice. Therefore, selective activation of KAT6A in PVIs of FC recapitulates the morphological impairment of PVIs and mouse behavioral alterations induced by CS, linking stress-responsive epigenetic changes in PVIs to cellular and behavioral outcomes.

## Discussion

Our study demonstrates that PVIs exhibit brain region- and experience-specific molecular plasticity, with distinct molecular programs engaged by CS and VR (**Supplementary Fig. 8**). Rather than representing opposing modulation of the same molecular program, CS and VR engage largely non-overlapping molecular responses, suggesting that positive and negative experiences may be associated with different regulatory programs in PVIs. At the functional level, these experience-dependent responses are organized into region-specific molecular architectures, suggesting that PVIs integrate environmental inputs through structured and context-dependent molecular programs rather than predominantly through a uniform bidirectional mechanism. In this context, *Kat6a* was identified as a CS-associated epigenetic regulator in PVIs and provides a mechanistic link between H3K23 acetylation-dependent chromatin remodeling and CS-associated structural and behavioral adaptations (**Supplementary Fig. 8**).

Previous studies on PVIs have mainly focused on its functional roles at the neural circuit and behavioral levels, such as the inhibition of excitatory neurons, regulation of network oscillations, and control of behavioral output^16,53^. In addition, PVIs are known to regulate critical-period plasticity during development and contribute to behavioral responses to antidepressant interventions through serotonin-dependent signaling pathways^54–56^. However, how PVIs respond to environmental challenges at the molecular level remain unclear. In this study, by examining cell-type-specific translational profiles, we discovered that PVIs exhibit significant molecular responses to environmental stimuli, and these responses depend on both the type of experience and the brain region where PVIs are embedded. Given that VR has been shown to mitigate CS-induced behavioral deficits^11,57^, one would expect that PVIs utilize the same gene network to respond to positive and negative experiences in opposing manners. Surprisingly, CS and VR recruit largely non-overlapping sets of gene networks even in the same brain region. This distinction is supported by co-expression network and pathway-level analyses, which consistently reveal that different molecular programs are associated with CS and VR. Together, these findings provide a conceptual framework for understanding how inhibitory neurons implement experience-dependent neural circuit remodeling at the molecular level and may offer a molecular basis for understanding how positive and negative experiences engage different molecular responses in PVIs. Therapeutic strategies aimed at alleviating CS-induced behavioral deficits may benefit from considering molecular pathways associated with positive experiences, rather than only focusing on reversing CS-associated changes.

VR-associated programs are enriched for gene sets related to synaptic function and cellular transport, whereas CS-associated programs more frequently involve structural, metabolic, and gene regulatory processes, as identified by GO and co-expression network analyses. At the pathway level, CS-associated responses showed a greater tendency toward cross-regional convergence, whereas VR-associated pathways were more region-specific. These patterns indicate that the organization of experience-dependent responses reflects region-dependent integration across brain regions. Pathways related to mitochondrial and metabolic processes were also represented within CS-associated clusters, consistent with previous findings implicating these processes in neurodevelopmental disorders^58,59^. These observations indicate that experience-dependent molecular adaptation is organized across multiple levels, from gene-level functional programs to higher-order pathway architectures, reinforcing the conclusion that positive and negative experiences are not encoded as opposing molecular states but instead engage distinct regulatory programs.

Among the genes exhibiting experience-dependent translational changes in PVIs, *Kat6a* showed consistent and FC-specific upregulation, which was not observed in hippocampal PVIs under the same conditions, indicating localized epigenetic regulation within frontal circuits. KAT6A is a member of the MYST family of histone acetyltransferases and has been implicated in neural progenitor self-renewal and early neural differentiation^60^. Pathogenic variants in KAT6A cause Arboleda-Tham syndrome, characterized by intellectual disability and developmental delay^61,62^. In mice, loss of Kat6a disrupts synaptic structure and plasticity in hippocampal CA3 neurons, leading to memory deficits^48^. These studies underscore the importance of KAT6A in neurodevelopment and synaptic regulation, yet its role in experience-dependent plasticity of inhibitory neurons has not been defined. Consistent with elevated KAT6A expression, CS increased acetylation at its substrate site H3K23 in FC PVIs and altered the genome-wide distribution of H3K23ac peaks. Functional enrichment analyses indicated that these chromatin changes preferentially affected genes involved in synaptic organization, neuronal signaling, and neuropsychiatric disease pathways, supporting a role for KAT6A-associated chromatin remodeling in shaping PVI responses to stress. Notably, CS-induced chromatin remodeling appeared to operate at a subgenic level, potentially reflecting differential regulation of promoters, enhancers, and gene bodies rather than uniform transcriptional activation or repression. This pattern suggests regulatory complexity in which chromatin modifications are redistributed within gene loci, enabling fine-tuned modulation of neuronal function under CS. In addition, genes associated with mixed acetylation patterns (both hyper- and hypo-acetylated peaks) showed stronger enrichment for AD related gene sets compared to genes with unidirectional acetylation changes, while all groups exhibited enrichment across multiple neuropsychiatric disease gene sets. In comparison, genes with unidirectional acetylation changes remained more strongly associated with canonical stress-response pathways. These observations suggest that locus-specific chromatin remodeling may define a distinct regulatory mechanism, in which mixed acetylation patterns are associated with pathways relevant to neuronal vulnerability and long-term dysfunction, rather than immediate stress-responsive processes. Functionally, targeted elevation of KAT6A in mPFC PVIs recapitulated key features of CS-associated structural and behavioral alterations consistent with CS exposure. The convergence of chromatin remodeling, morphological changes, and behavioral deficits suggests a link between CS-induced epigenetic regulation and inhibitory circuit dysfunction and provides functional support for the role of KAT6A in mediating CS-induced plasticity in PVIs. Together, our findings position KAT6A as a region-specific CS-responsive epigenetic regulator associated with molecular and morphological remodeling in PVIs.

A major challenge in gene-environment interaction research is to define how environmental exposures are translated into cell-type-specific and circuit-level molecular programs. While recent studies have begun to profile molecular adaptations to social experience, stress, and exercise using more refined or multimodal approaches^33–36^, these efforts have not directly compared opposing experiences within the same inhibitory cell type across regions. Our PVI-specific translatome and network analyses address this gap by showing that opposing experiences, CS and VR, are organized into structured, region-dependent molecular programs within PVIs rather than a single bidirectional mechanism. This framework is supported by translational profiling, which captures ribosome-associated transcripts that may be underrepresented in nucleus-based sequencing approaches, particularly in highly polarized neurons such as PVIs, where mRNAs are distributed across dendritic and axonal compartments^63–65^. This cell-type-specific organization provides a molecular basis for understanding how environmental exposures may interact with genetic vulnerability. In our dataset, CS-responsive DTGs and genes linked to H3K23 acetylation changes in FC PVIs were significantly enriched for psychiatric risk-associated gene sets, including SCZ, ASD, BD, and depression. A recent study identified enrichment of ASD-, SCZ-, and BD-associated genes in PVIs within the hippocampal regions in response to social simulation and discovered molecular programs linked to experience-dependent PVI plasticity^36^. Together, these findings suggest that experience-driven remodeling of inhibitory neurons can converge on established risk networks, offering a potential mechanistic entry point for how environmental factors may modulate disease susceptibility.

In summary, our study demonstrates that opposing environmental experiences engage PVIs through structured, region-dependent molecular programs rather than a uniform bidirectional mechanism. We reveal how environmental exposures are translated into molecular programs within inhibitory neurons. By defining this cell-type-specific architecture and identifying KAT6A as a CS-responsive epigenetic regulator associated with molecular and structural changes in PVIs, we highlight a potential epigenetic pathway linking CS exposure to inhibitory circuit remodeling. These findings offer a molecular perspective on how gene-environment interactions may shape circuit organization and interact with molecular pathways implicated in neuropsychiatric disorders.

### Limitation of this study

We recognize that a limitation of our study is that, while translational profiling has its advantage in capturing mature mRNA transcripts throughout the complex neuronal cell bodies, processes, and terminals, it does not resolve molecular heterogeneity among PV interneuron subtypes. In addition, MERFISH is limited to assessing mRNA levels, which is not informative to assess translational changes. Future translational profiling at single cell levels, especially with spatial resolution, will be important to resolve experience induced gene network changes in distinct PVI subtypes. Another consideration is that the degree of overlap between DTG sets may be influenced by the analytical framework used for differential translational analyses. Future studies using complementary analytical approaches may help further refine estimates of shared and experience-specific molecular responses in PVIs. In addition, we observed discrepancies between RiboTag-seq and IHC results for a subset of candidate genes. Such differences could be due to the fact that RiboTag-seq captures ribosome-associated mRNA whereas IHC reflects accumulated protein levels. Since in neurons, translational activity, protein stability, and subcellular localization are often uncoupled, the differences between translational and protein-level measurements may provide opportunities to investigate how these molecular processes are regulated in neurons.

## Methods

Detailed methods are provided in the supplemental file for review.

## Data availability

Raw and processed RiboTag-seq and CUT&Tag data have been deposited in the NCBI Gene Expression Omnibus (GEO) under accession numbers GSE334645 (RiboTag-seq) and GSE334647 (CUT&Tag-seq), respectively.

## Supplemental information

There are 6 supplemental figures (see Supplemental Figures and Methods file) and 7 supplemental tables.

## Acknowledgments

We thank I. Dhawan, E. Chang, T. Korabelnikov, N. Méndez-Albelo, S. Kannan, Y. Xing in the Zhao lab for technical assistance; P. Kumarage and S. Liu at Waisman Center Data Science Core for MERFISH analysis, J. Panksepp, M. Eastwood, and K. Knobel at the Waisman IDD Model Core for core services; Dr. A. M. M. Sousa for input on manuscript. This work was supported by grants from the National Institutes of Health (R01NS105200, R01MH116582, R01MH118827, R01MH136152, R01NS138268 to X.Z.; 5R44GM151926 and American Heart Association 26RIRA1649090 to A.H.R.; P50HD105353 to Waisman Center; R21NS142497 to Y.Gao), DOD IIRA grant W81XWH-22-1-0621 (to X.Z.), Brain Research Foundation Scientific Innovation Award, SFARI pilot grant, Eagles Autism Foundation pilot grant, Vilas Mid-Career Award, Jenni and Kyle Professorship, Kellet Mid-Career Award, and Vilas Distinguished Achievement Professorship (to X.Z.), Hilldale Undergraduate fellowship (to S.X.H. and E.D.J.) Morris Aprison Scholarship for Undergraduate Research (to S.X.H. and P.Y.), postdoctoral/predoctoral fellowships from FRAXA (to M.J., Z.X. and J.Lee), and Warren Alpert Distinguished Scholarship (to Y. Guo).

## Author contributions

X.Z. conceived the concept, designed experiments and supervised the project. Y.Guo designed and performed experiments, collected and analyzed data, interpreted results, prepared figures. Y.Guo. and X.Z. wrote the manuscript. Y.Gao conceived the original project concept, initiated the project, generated the RiboTag-seq dataset, and contributed to MERFISH experiments. S.K. performed bioinformatics analysis. Z.X. and M.S. performed behavior tests, Y.Z. performed the GSVA analysis. E.K. and A.R. provided guidance for nuclear isolation and CUT&Tag optimization. J.Lee., P.Y., S.X.H, E.D.J, D.N.T., J.T.W., J.Le. and K.A.S. collected data.

