## Supplementary figures for "Opposing Experiences Shape Parvalbumin Interneuron Plasticity through Distinct Molecular Programs"

**Supplementary Figures and Methods**

### Opposing experiences shape parvalbumin interneuron plasticity through distinct and brain region-specific molecular programs

*Guo et al, 2026*

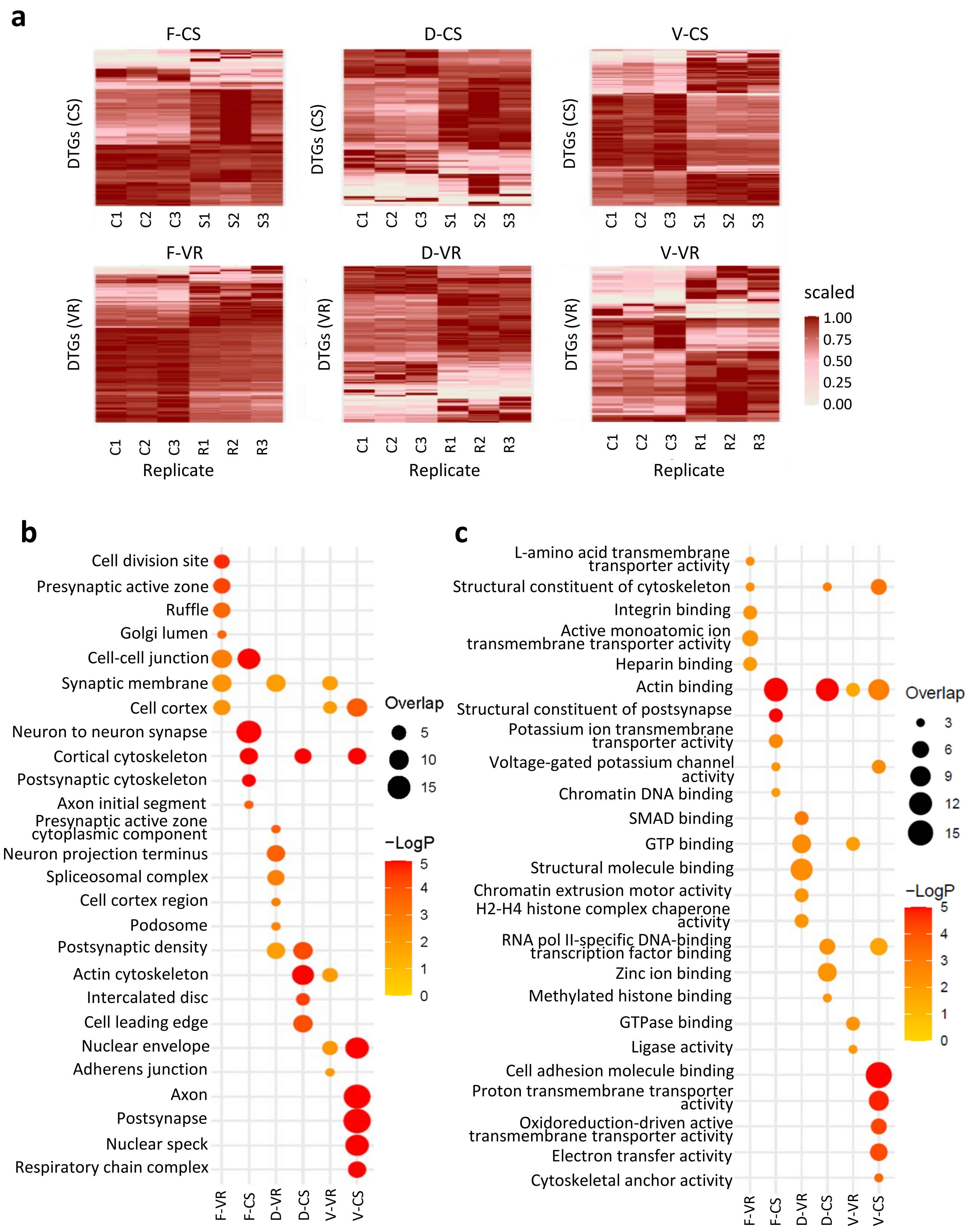
a
F-CS
D-CS
V-CS
DTGs (CS)
C1 C2 C3 S1 S2 S3
F-VR
D-VR
V-VR
DTGs (VR)
C1 C2 C3 R1 R2 R3
Replicate
scaled
1.00
0.75
0.50
0.25
0.00
b
Cell division site
Presynaptic active zone
Ruffle
Golgi lumen
Cell-cell junction
Synaptic membrane
Cell cortex
Neuron to neuron synapse
Cortical cytoskeleton
Postsynaptic cytoskeleton
Axon initial segment
Presynaptic active zone cytoplasmic component
Neuron projection terminus
Spliceosomal complex
Cell cortex region
Podosome
Postsynaptic density
Actin cytoskeleton
Intercalated disc
Cell leading edge
Nuclear envelope
Adherens junction
Axon
Postsynapse
Nuclear speck
Respiratory chain complex
F-VR F-CS D-VR D-CS V-VR V-CS
Overlap
5
10
15
−LogP
5
4
3
2
1
0
c
L-amino acid transmembrane transporter activity
Structural constituent of cytoskeleton
Integrin binding
Active monoatomic ion transmembrane transporter activity
Heparin binding
Actin binding
Structural constituent of postsynapse
Potassium ion transmembrane transporter activity
Voltage-gated potassium channel activity
Chromatin DNA binding
SMAD binding
GTP binding
Structural molecule binding
Chromatin extrusion motor activity
H2-H4 histone complex chaperone activity
RNA pol II-specific DNA-binding transcription factor binding
Zinc ion binding
Methylated histone binding
GTPase binding
Ligase activity
Cell adhesion molecule binding
Proton transmembrane transporter activity
Oxidoreduction-driven active transmembrane transporter activity
Electron transfer activity
Cytoskeletal anchor activity
F-VR F-CS D-VR D-CS V-VR V-CS
Overlap
3
6
9
12
15
−LogP
5
4
3
2
1
0

**Supplementary Fig. 1 | Heatmaps and GO enrichment analyses of differentially translated genes (DTGs) in PVIs under CS and VR. a,** Heatmaps of differentially translated mRNAs in PVIs from Frontal cortex, dorsal hippocampus and ventral hippocampus under CS or VR compared to control. Each column represents a biological replicate. Color scale indicates scaled median-normalized expression values (0–1) derived from the EBSeq analysis (0–1), with darker red reflecting higher relative expression. **b,c,** Top five enriched GO Cellular Component (CC, **b**) and Molecular Function (MF, **c**) terms for each DTG set were selected, and significant enrichments ($p < 0.05$) for these terms were displayed across all six DTG sets. CS, S: chronic stress; Ctrl, C: control; VR, R: voluntary running. F: Frontal cortex; D: dorsal hippocampus; V: ventral hippocampus.

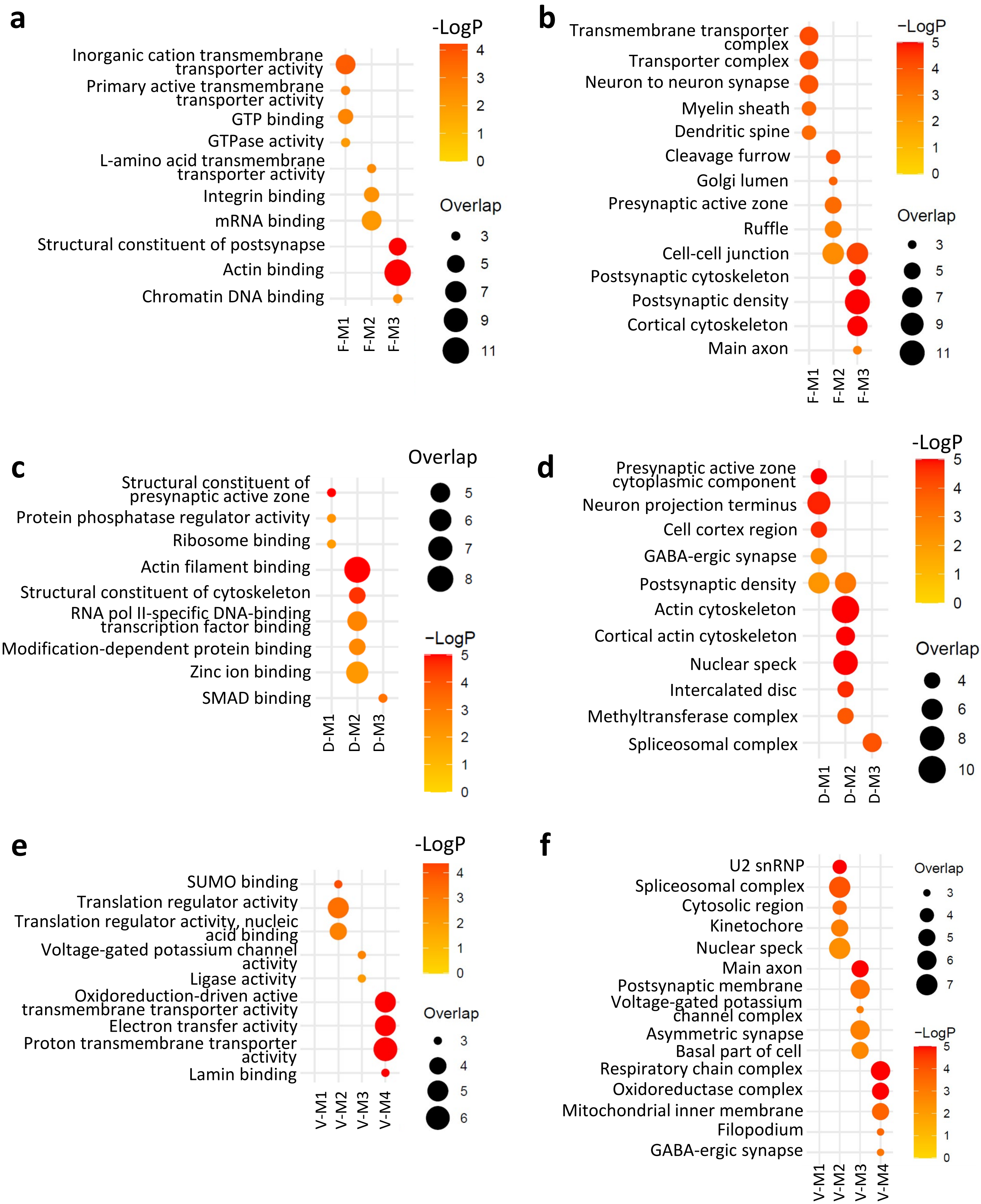
a
-LogP
Inorganic cation transmembrane transporter activity
Primary active transmembrane transporter activity
GTP binding
GTPase activity
L-amino acid transmembrane transporter activity
Integrin binding
mRNA binding
Structural constituent of postsynapse
Actin binding
Chromatin DNA binding
F-M1
F-M2
F-M3
Overlap
b
Transmembrane transporter complex
Transporter complex
Neuron to neuron synapse
Myelin sheath
Dendritic spine
Cleavage furrow
Golgi lumen
Presynaptic active zone
Ruffle
Cell-cell junction
Postsynaptic cytoskeleton
Postsynaptic density
Cortical cytoskeleton
Main axon
F-M1
F-M2
F-M3
−LogP
Overlap
c
Structural constituent of presynaptic active zone
Protein phosphatase regulator activity
Ribosome binding
Actin filament binding
Structural constituent of cytoskeleton
RNA pol II-specific DNA-binding transcription factor binding
Modification-dependent protein binding
Zinc ion binding
SMAD binding
D-M1
D-M2
D-M3
Overlap
−LogP
d
Presynaptic active zone cytoplasmic component
Neuron projection terminus
Cell cortex region
GABA-ergic synapse
Postsynaptic density
Actin cytoskeleton
Cortical actin cytoskeleton
Nuclear speck
Intercalated disc
Methyltransferase complex
Spliceosomal complex
D-M1
D-M2
D-M3
-LogP
Overlap
e
SUMO binding
Translation regulator activity
Translation regulator activity, nucleic acid binding
Voltage-gated potassium channel activity
Ligase activity
Oxidoreduction-driven active transmembrane transporter activity
Electron transfer activity
Proton transmembrane transporter activity
Lamin binding
V-M1
V-M2
V-M3
V-M4
-LogP
Overlap
f
U2 snRNP
Spliceosomal complex
Cytosolic region
Kinetochore
Nuclear speck
Main axon
Postsynaptic membrane
Voltage-gated potassium channel complex
Asymmetric synapse
Basal part of cell
Respiratory chain complex
Oxidoreductase complex
Mitochondrial inner membrane
Filopodium
GABA-ergic synapse
V-M1
V-M2
V-M3
V-M4
Overlap
−LogP

**Supplementary Fig. 2 | GO enrichment analysis of DTGs within each WGCNA module. a,b,** Representative top GOCC (**a**) and GOMF (**b**) terms are shown for each FC module. **c,d,** Representative top GOCC (**c**) and GOMF (**d**) terms are shown for each dHC module. **e,f,** Representative top GOCC (**e**) and GOMF (**f**) terms are shown for each vHC module. The top five most significant terms for each module DT gene set were selected and the significant enrichments ($p<0.05$) of these terms were plotted for across all modules.

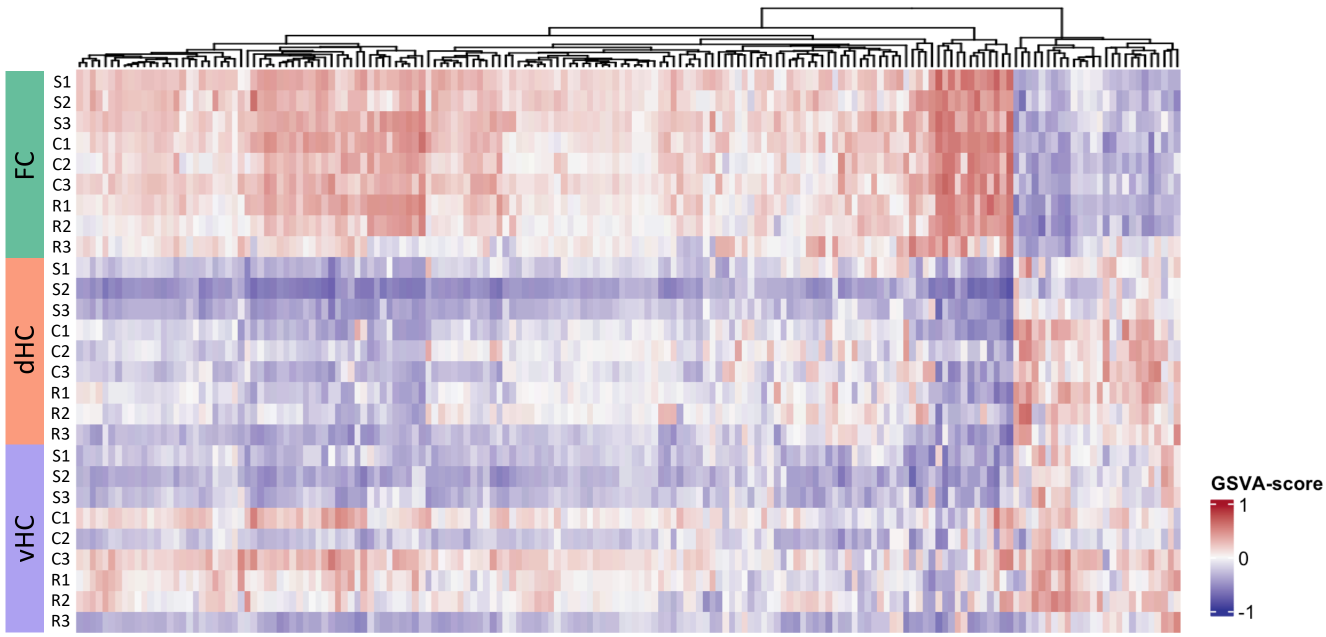


**Supplementary Fig. 3 | Gene set variation analysis (GSVA) of region- and experience-specific pathway activity in PVIs.** GSVA heatmap of selected neuronal-intrinsic (development, maturation, and migration) and behavior-associated terms, showing stronger positive enrichment in the FC compared with dHC and vHC. Color intensity represents relative GSVA enrichment score per sample, with warmer colors indicating higher pathway activity.

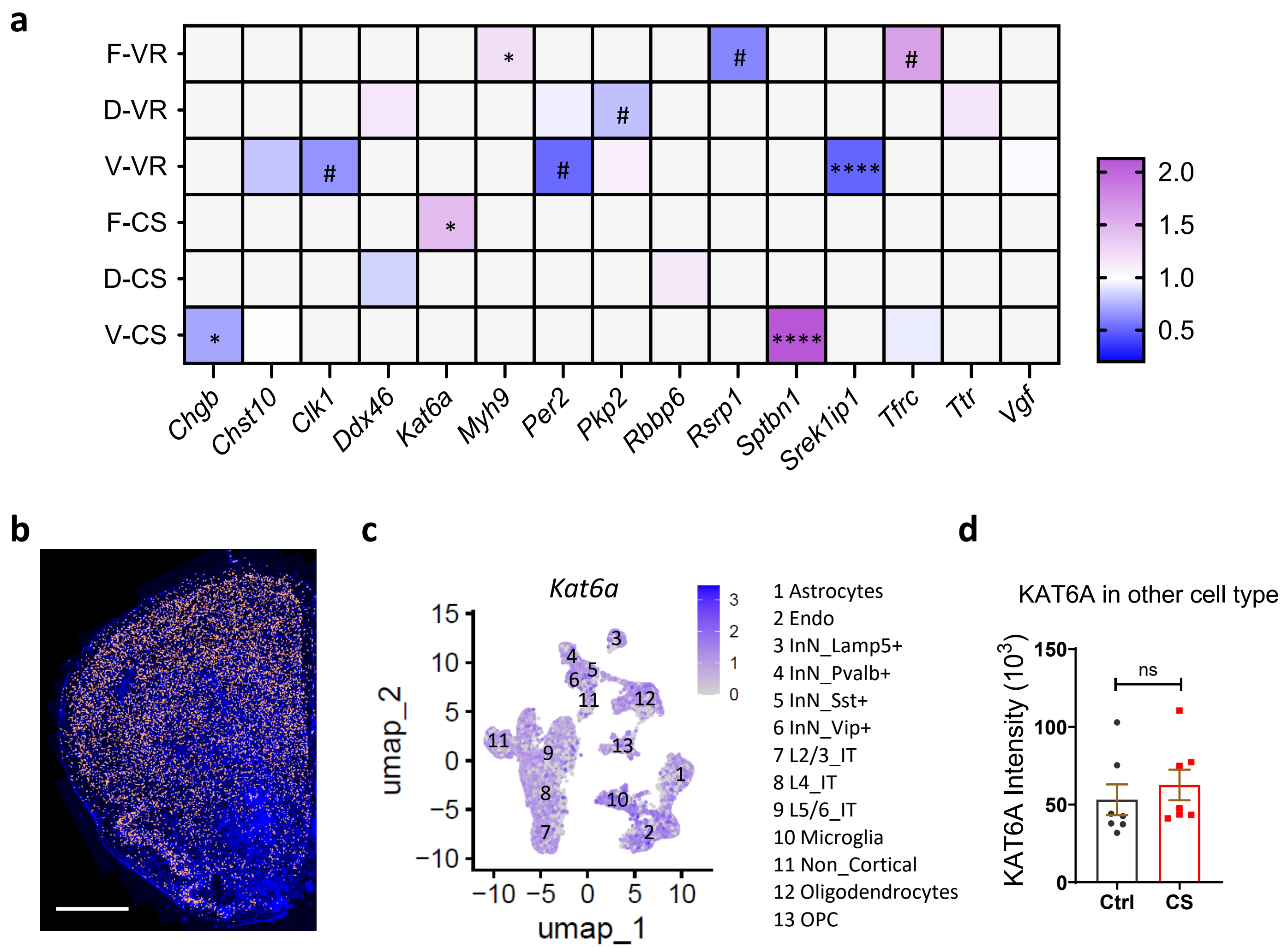


**Supplementary Fig. 4 | Validation of candidate genes and cell-type-specific upregulation of KAT6A in FC-PVIs. a,** Summary of IHC validation results. * indicate statistically significant differences consistent with RiboTag-seq, while # indicate significant differences in the opposite direction. **b,** Merscope image showing the location of *Kat6a* transcripts in mouse brain section. Scale bar: 1mm. **c,** Feature plot showing the expression level of *Kat6a* transcripts across all cell types. **d,** Quantification demonstrating that KAT6A expression was specifically increased in the FC-PVIs under CS. Data are presented as mean ± s.e.m from N ≥ 3 mice. Two-tailed Student's test in (**a**) and (**d**). *$p < 0.05$, ****$p < 0.0001$.

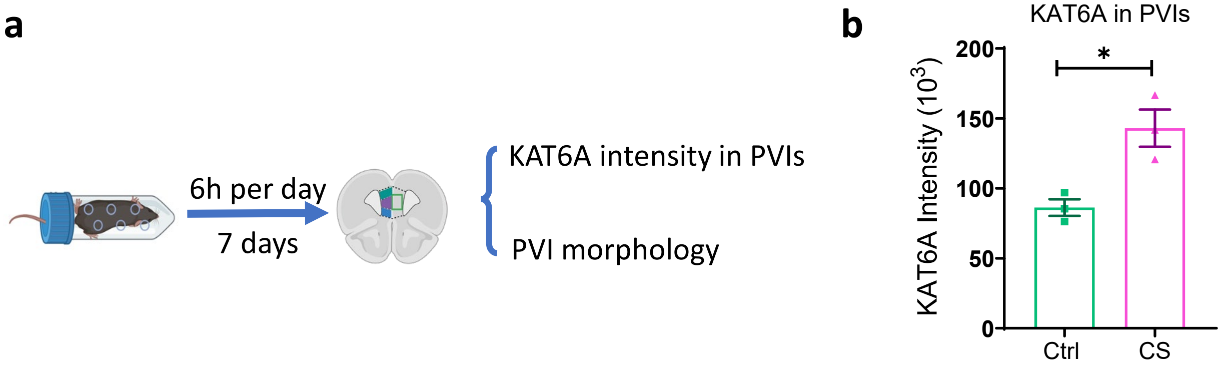


**Supplementary Fig. 5 | Validation of the revised stress paradigm. a,** Experimental scheme for revised chronic stress (CS) paradigm used for PVI nuclear isolation, CUT&Tag profiling, and subsequent behavioral studies. **b,** Quantification of KAT6A intensity in mPFC PVIs from mice subjected to the revised CS paradigm shown in (**a**), validating KAT6A upregulation under the modified stress condition. Data are presented as mean ± s.e.m from N = 3 mice, two-tailed Student's test, $*p < 0.05$.

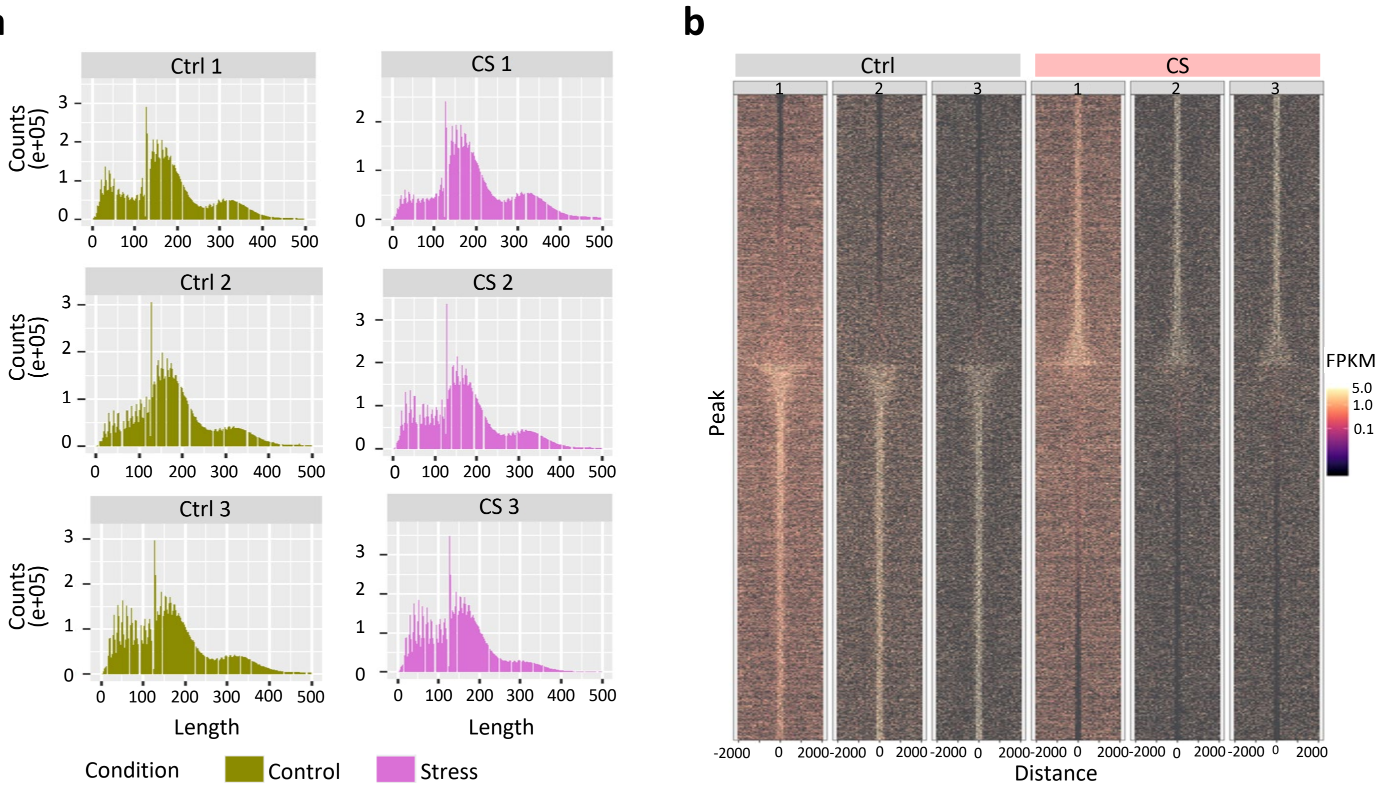


c

| Rank | Top enriched Motifs | Name | −log10(p) | q-value (Benjamini) | % of Targets Sequences |
|---|---|---|---|---|---|
| 1 | 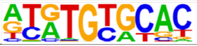 | Zscan4c | 7.209 | 0.3492 | 7.16% |
| 2 | 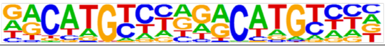 | p53 | 6.669 | 0.3492 | 1.69% |
| 3 | 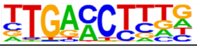 | RARa(NR) | 5.866 | 0.3492 | 32.20% |
| 4 | 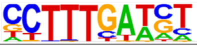 | LEF1(HMG) | 4.958 | 0.6631 | 7.60% |
| 5 | 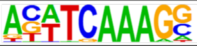 | Tcf3(HMG) | 4.830 | 0.6631 | 3.02% |

d

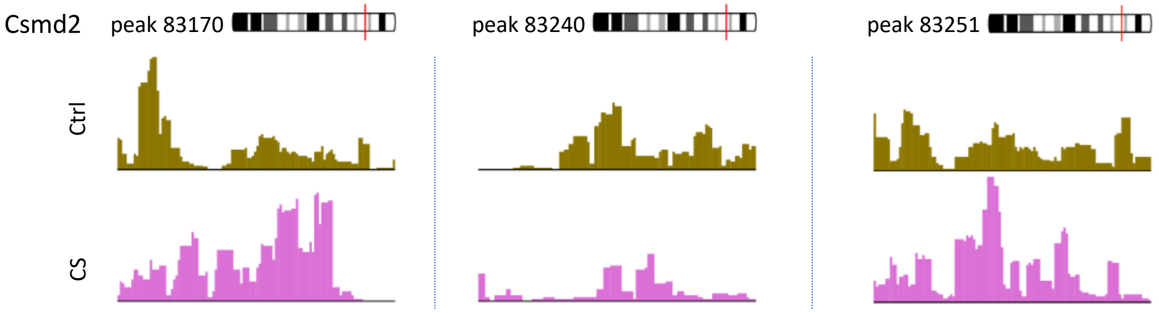

**Supplementary Fig. 6 | Additional CUT&Tag analyses. a,** Fragment length distribution of Ctrl and CS CUT&Tag libraries. Peaks at ~150 bp and ~300 bp correspond to mono- and di-nucleosome fragments, respectively. **b,** Signal intensity plots for individual biological replicate. **c,** Motif enrichment analysis of H3K23ac DA regions identified between CS and Ctrl samples. **d,** Peak visualization of H3K23ac profiles at *Csmd2 locus*, showing concurrent hyper- and hypo-acetylation at distinct regions within the same gene under CS compared with Ctrl.

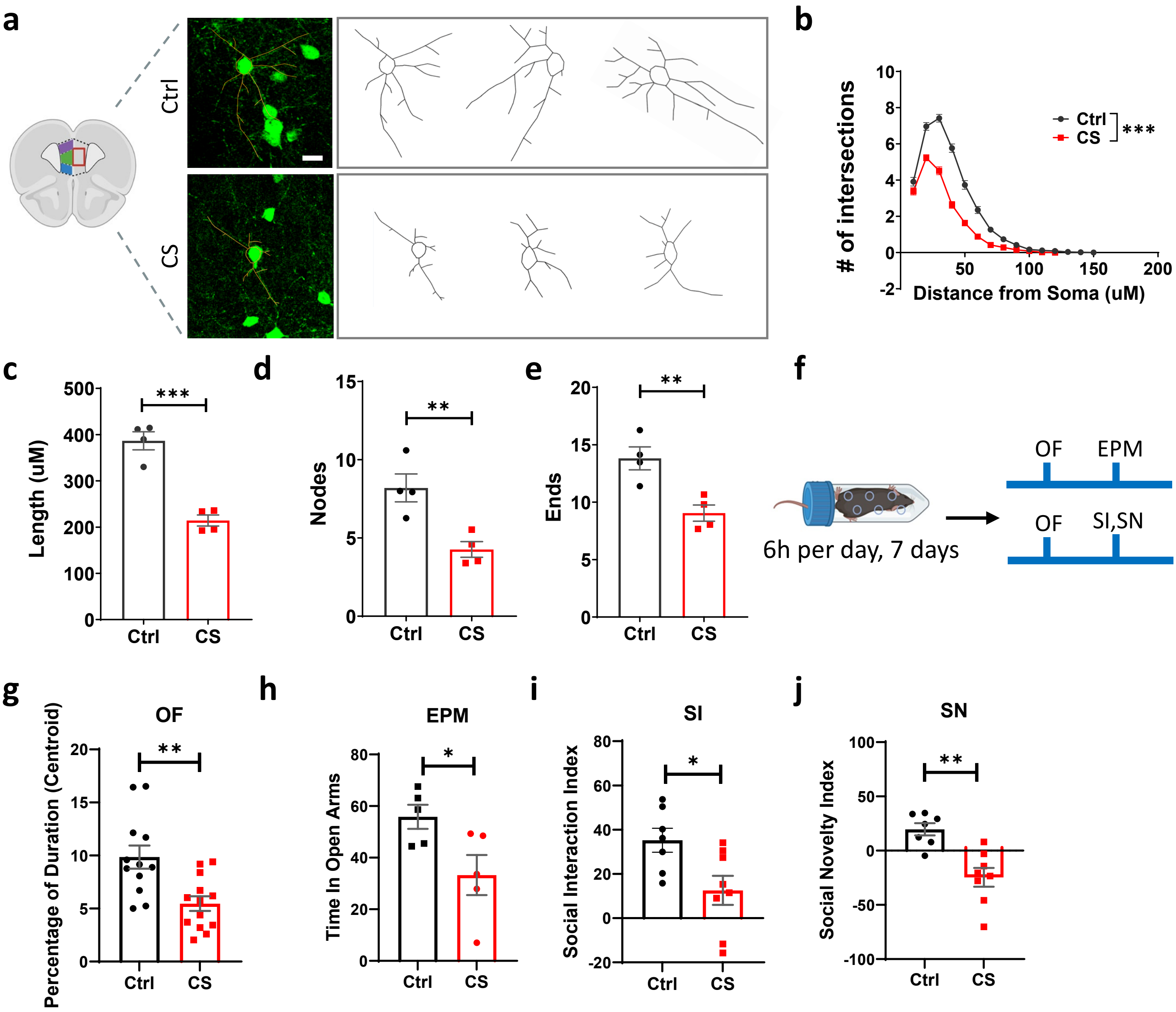


**Supplementary Fig. 7 | CS leads to reduced PVI complexity and behavior deficits in wt mouse. a,** Representative confocal images and traces of PV+ neurons for assessing the effect of stress on neuronal morphology. Scale bar, 20 μm. **b,** Sholl analysis of PVIs in CS and Ctrl mice. **c-e**, Total neurite length (**c**), total number of nodes (**d**) and total neurite endings (**e**). **f,** Experimental diagram of behavior tests for CS mice. **g,h,** Anxiety levels assessed by open field (OF, **g**) test and elevated plus maze (EPM, **h**) test. **i,j,** CS mice exhibited reduced social interest (SI, **i**), and social recognition (SN, **j**). Data are presented as mean ± s.e.m. MANOVA in (**b**), $F(1,58) = 132.643$, $p < 0.001$, n = 60 cells from 4 mice. Two-tailed Student's test in (**c-e**) and (**g-j**) from N ≥ 4 mice. *p < 0.05, **p < 0.01, ***p < 0.001.

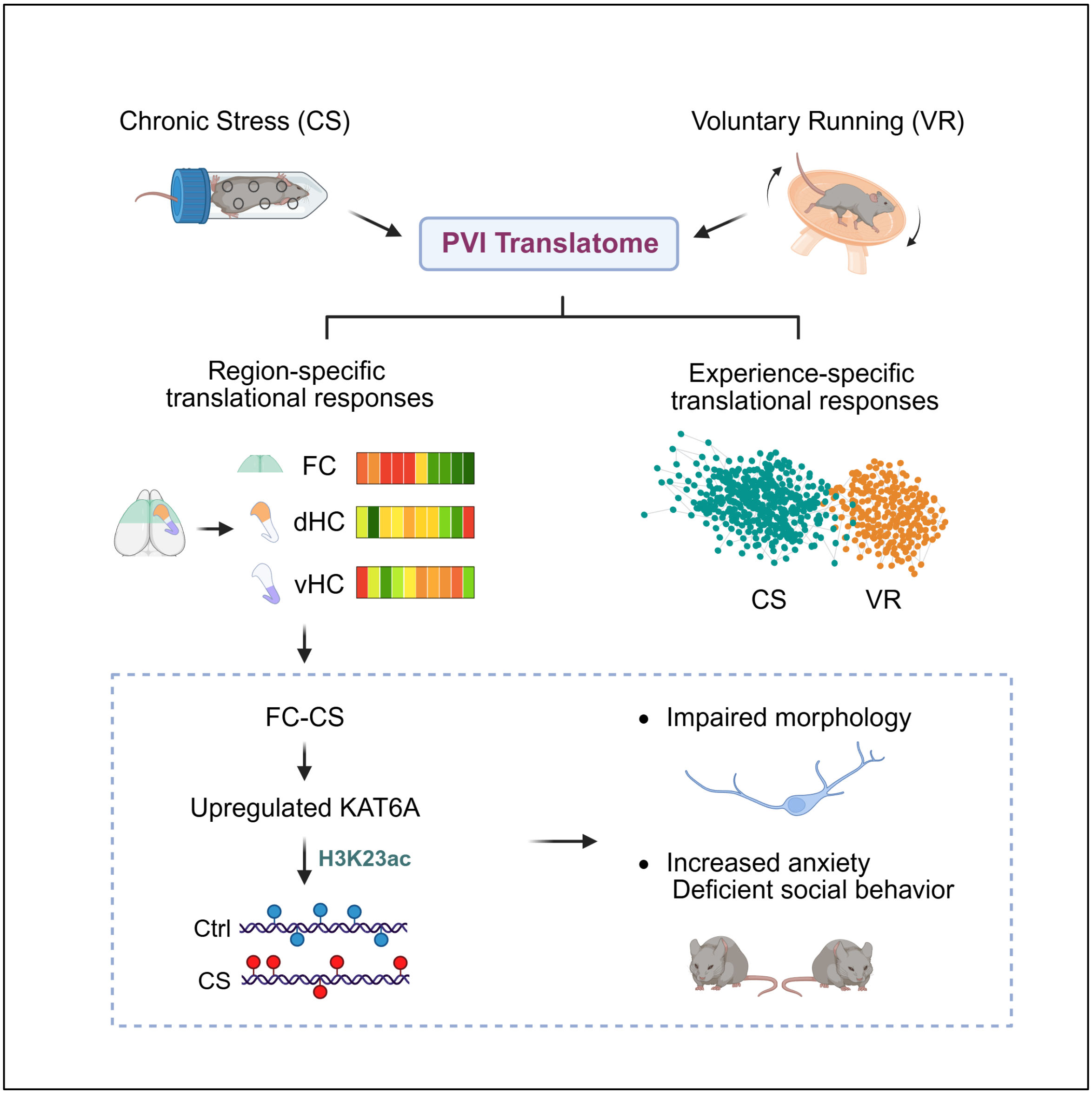


**Supplementary Fig. 8 | Schematic summary of region- and experience-specific PVI responses and the effects of chronic stress on PVI morphology and behavior**. PVIs exhibit brain region- and experience-specific molecular plasticity, with distinct molecular programs engaged by chronic stress (CS) and voluntary running (VR). Rather than oppositely modulating a shared molecular program, CS and VR elicit largely non-overlapping molecular responses, suggesting distinct regulatory programs underlying PVI adaptations to positive and negative experiences. Kat6a was identified as a CS-associated epigenetic regulator in PVIs, linking H3K23 acetylation to CS-associated changes in PVI dendritic morphology and behavior.
